## Supplemental Material for "High-resolution Nanopore methylome-maps reveal random hyper-methylation at CpG-poor regions as driver of chemoresistance in leukemias"

[1]Department of Information Engineering, University of Florence, Florence, Italy,. [2]Institute for Biomedical Technologies, National Research Council, Segrate, Milano, Italy. [3]Department of Experimental and Clinical Medicine, University of Florence, Florence, Italy. [4]Department of Experimental Oncology, IEO European Institute of Oncology IRCCS, Milano, Italy. [5]Department of Oncology and Hemato-Oncology, University of Milan, Milan, Italy. [6]Azienda Socio-Sanitaria Territoriale Papa Giovanni XXIII, Bergamo, Italy. [7]Clinica Ematologica, Azienda Sanitaria Universitaria Integrata di Udine, Udine, Italy. [*] **Correspondence**: Alberto Magi, Department of Information Engineering, University of Florence, 50100, Florence, Italy, Pier Giuseppe Pelicci, Department of Experimental Oncology, IEO European Institute of Oncology IRCCS, Milano, Italy . [+] These authors contributed equally to this work.

### Nanopore library preparation and sequencing

DNA was qualitatively and quantitatively analyzed using Nanodrop and Qubit respectively. The sequencing was performed on the GridION 5X platform (Oxford Nanopore Technologies, Oxford, UK). Library preparation was performed according to the Oxford Nanopore Technologies (ONT) manufacturer's protocol for genomic DNA, the Ligation Sequencing Kit 1D SQK-LSK109. For each library 1 ug of DNA as starting material was used. The input DNA for all nanopore libraries was unshared. Initially DNA was mixed with 3,5 ul of NEBNext FFPE DNA Repair Buffer and 2 ul of NEBNext FFPE DNA Repair Mix (M6630, NEB, Ipswich, Massachusetts) in order to repair possible nicks, and then end-prepped adding 3,5 ul of Ultra II End-prep reaction buffer and 3 ul of Ultra II End-prep enzyme mix (E7546, NEB). The reaction was incubated at 20°C for 5 minutes and 65°C for 5 minutes, followed by a purification step with 60 ul of 1X AMPure XP beads, by a washing and an elution. The end-repaired DNA was ligated with 5 ul Adapter Mix (AMX, ONT) using 10 ul NEBNext Quick T4 DNA Ligase (E6056, NEB) for 10 min at room temperature. The adapted-ligated DNA was cleaned up by adding 40 ul of 1X AMPure XP beads and incubated at room temperature for 5 min. The beads were pelleted on a magnetic rack and pellet was washed twice by resuspending in 250 ul Long Fragment Buffer (LFB, ONT) in order to enrich for long molecules. Then, the reaction was eluted in 15 ul of Eluition Buffer (EB, ONT) at room temperature for 10 minutes. The prepared library was quantified by Qubit and then, prior to loading onto the flow cell, was mixed with 37,5 ul of Sequencing Buffer (SB, ONT) and 25,5 ul of Loading Beads (LB, ONT). According to the ONT's protocol before loading, each R9.4.1 SpotON flow cell (FLO-MIN106D) was primed mixing the Flush Tether (FLT, ONT) directly to the tube of Flush Buffer (FB, ONT) and loading this mix via the priming port. Libraries were sequenced on a GridION 5x device and, in order to maximize the total throughput, sequencing was carried out for 72 h.

### Shifting Level Models with truncated gaussian

Classical SLM are a class of models (Magi *et al.*, 2010) in which sequential observations $x = (x_1, \ldots, x_i, \ldots, x_N)$ are considered to be realizations of the sum of two independent stochastic processes

$$x_i = m_i + \epsilon_i,$$

$$m_i = (1 - z_{i-1}) \cdot m_{i-1} + z_{i-1} \cdot (\mu + \delta_i)$$

where $m_i$ is the unobserved mean level and $\epsilon_i$ is normally distributed white noise. The process $m_i$ changes its value independently of $m_{i-1}$ and is controlled by the process $z_i$: when $z_{i-1} = 0$ , $m_i$ is the same as $m_{i-1}$ and when $z_{i-1} = 1$, $m_i$ is incremented by the normal random variable $\delta_i$ ($\delta_i \sim N(0, \sigma_\mu^2)$). $z_1, z_2, ...$ are independent and identically distributed random variables taking the values 0,1 with probabilities $\eta = Pr(z_i = 1)$, $1 - \eta = Pr(z_i = 0)$.

To take into account the sparse distribution of CpGs along the genome, as in (Magi *et al.*, 2013) we extended the classical SLM to an heterogeneous form, where the probability $Pr(z_i = 1)$ depends on this distance between consecutive CpGs ($d_i$) with the following formula:

$$\Pr(\mathrm{z}_i = 1) = \eta(d_i) = \frac{1}{2} \cdot \theta + ((\frac{1}{2} - \theta) \cdot exp\left[\frac{log(\theta)}{\frac{d_i}{d_{Norm}}}\right]) \ (1)$$

where $\eta(d_i)$ is the probability of random variables $z_i$ to be equal to 1, $\theta$ is a constant parameter , $d_i$ is the distance between the $i - th$ and $i_1 - th$ CpG and $d_{Norm}$ is the distance normalization parameter. Equation 1 defines the dependence between the probability $Pr(z_i = 1)$ and the genomic distance between adjacent CpGs $d_i$: the larger is $d_i$ and the larger is $Pr(z_i = 1)$ and consequently the larger is the probability to jump between two mean levels $m_i$. The constant parameter $\theta$ can be seen as the baseline probability of random variables $z_i$ to take value 1 while the $d_{Norm}$ parameter modulates the genomic distance at which the probability $Pr(z_i = 1)$ begins to grow: for distances smaller than $d_{Norm}$ the probability $Pr(z_i = 1) = (\theta - \theta^2) \sim \theta$, while when $d_i$ is larger than $d_{Norm}$ the probability $Pr(z_i = 1)$ grows until reaching the value 1.

The expected value of $x_i$ is equal to $\mu$ and, since the two stochastic processes are independent, the variance of $x_i$ is the sum of the variances of the two processes:

$$E[x_i] = \mu, \tag{2}$$

$$var[x_i] = \sigma_\mu^2 + \sigma_\epsilon^2. \tag{3}$$

In this way, SLM allows one to break up the total variance of the genomic profile in two parts: the biological variance ($\sigma_\mu^2$) and the experimental variance ($\sigma_\epsilon^2$). Using (3) we can introduce a different parametrization of the SLM by defining the parameter $\omega = \sigma_\mu^2/\sigma^2$ (with $\sigma^2 = var[x_i]$) such that $\sigma_\mu^2$ by $\omega \cdot \sigma^2$ and $\sigma_\epsilon^2$ by $(1 - \omega)\sigma^2$.

With this new parametrization the joint probability distribution of the observations and latent variables $p(x, m, z|\Theta)$ , given the parameters, can be explicated in the following:

$$\begin{aligned} &p(x, m, z|\theta) = p(x|m, \sigma^2, \omega) \cdot p(m|z, \mu, \sigma^2, \omega) \cdot p(z|\eta(d_i)) = \\ &= \prod_{i=1}^{N} p(x_i|m_i, \sigma^2, \omega) \cdot p(m_0) \times \prod_{i=0}^{N} p(m_{i+1}|m_i, z_i, \mu, \sigma^2, \omega) \cdot p(z_i|\eta(d_i)), \end{aligned} \tag{4}$$

Equation 4 defines an Heterogeneous Hidden Markov Model (HHMM) of order one, in which a single state variable, $q_i = (m_i, z_i)$, summarizes all the relevant past information of the underlying process.
In the model defined by (4) the elements of the HHMM are the following:

- the state transition probability distribution is:
  $p(q_{i+1}|q_i, \theta) = p(m_{i+1}|m_i, z_i, \mu, \sigma^2, \omega) \cdot p(z_i|\eta(d_i))$
- the emission probability distribution is:
  $p(x_i|q_i, \theta) = p(x_i|m_i, \sigma^2, \omega)$
- the initial state probability distribution is:
  $p(q_0|\theta) = p(m_0|\mu, \sigma^2, \omega)$

Since $\Delta\beta$ takes values in the range [-1,1], we modeled $\epsilon_i$ with a truncated gaussian distributions with upper and lower bound 1 and -1 respectively.

The fact that SLM is an HMM allows us to make use of the several algorithms developed for these kinds of models. To handle the infinite dimensionality of SLM, we use the same approach we previously developed in (Magi *et al.*, 2010).
We introduce a Markovian stochastic process $s_1, s_2, ..., s_k$ taking values in $S = \{1, 2, ..., K\}$. From equation 4, we know that the probability distribution of $x_i$, given the parameters, has the following form:

$$p(x_i|m_i, \sigma^2, \omega) = N(x_i|m_i, (1-\omega) \cdot \sigma^2), \tag{5}$$

Since $\Delta\beta$ takes values in the range [-1,1], we assume that the conditional probability of $x_i$, given $s_i = k$ is a truncated gaussian distributions with upper and lower bound 1 and -1 respectively, $N_{-1}^{1}(\mu_k, \sigma_\epsilon^2)$.

The parameter $\mu_k (k = 1, 2, ..., K)$ is associated to each state of the Markovian stochastic process and represents an approximation of the $m_i$ latent variables of the SLM.
The emission function of the HMM is defined as follows:

$$f_k(x) = \frac{1}{\sqrt{2\pi}\sigma_\epsilon} \exp\left[-\frac{1}{2}\left(\frac{x-\mu_k}{\sigma_\epsilon}\right)^2\right] \cdot \frac{1}{\Phi(\frac{1-\mu_k}{\sigma_\epsilon}) - \Phi(\frac{-1-\mu_k}{\sigma_\epsilon})}. \tag{6}$$

where $\Phi$ represent the cumulative distribution functions of a non-truncated Gaussian of parameter $\mu_k$ and $\sigma_\epsilon$.

To complete the description of the model, it remains to specify the state transition matrix $P$. From (4) the state transition probability has the following form:

$$\begin{aligned} p(m_{i+1}|m_i, z_i, \mu, \sigma^2, \omega) \cdot p(z_i|\eta(d_i)) &= [(1-z_i) \cdot \delta(m_{i+1} - m_i) + z_i \cdot N(m_{i+1}|\mu, \omega \cdot \sigma^2)] \cdot \\ &\quad \cdot [\eta(d_i) \cdot \delta(z_i - 1) + (1 - \eta(d_i)) \cdot \delta(z_i)], \end{aligned} \tag{7}$$

hence the state transition matrix is:

$$P_{jk} = \begin{cases} (1-\eta(d_i)) + \eta(d_i) \cdot g_{jk} & j = k \\ \eta(d_i) \cdot g_{jk} & j \neq k \end{cases} \tag{8}$$

where

$$\begin{aligned} g_{jk} &= c_j \cdot e^{-\frac{(\mu_k-\mu)^2}{2\sigma_\mu^2}}, \\ c_j &= \left( \sum_{k=1}^{K} e^{-\frac{(\mu_k-\mu)^2}{2\sigma_\mu^2}} \right)^{-1}. \end{aligned} \tag{9}$$

To estimate the parameters of the Heterogeneous truncated-gaussian Shifting Level Model (HT-GSLM), we develop a two-step algorithm that follows our previous idea in Magi *et al.* (2013). Since the$\Delta\beta$ can take values in a well-defined range, we used a large number of states K0 and we choose $\mu_k$ in order to densely and homogeneously cover the range [-1,1] instead of estimating the $\mu_k$ parameters by using the Baum and Welch algorithm (Magi *et al.*, 2010). Moreover, since we expect that the great majority of CpG have no differential methylation between samples ($\Delta\beta \sim 0$), we initialize $\mu = 0$. This simple solution drastically improves the computational performance of our algorithm without affecting its accuracy in the detection of signal shifts.

In the first step of the algorithm we initialize the mean $\mu$ and the variances $\sigma^2$, $\sigma_\mu^2$ and $\sigma_\epsilon^2$ with the following formulas:

$$\begin{aligned} &\mu = 0, \\ &\sigma = \sqrt{\frac{\sum_{i=1}^{N}(x_i-\mu)^2}{(N-1)}}, \\ &\sigma_\mu^2 = \omega \cdot \sigma^2, \\ &\sigma_\epsilon^2 = (1-\omega) \cdot \sigma^2. \end{aligned} \tag{10}$$

In the second step we apply the Viterbi algorithm to find the best state sequence $s^{(j)}$ and estimate the points of mean shift $z_i$. After Viterbi algorithm we calculate the median of the $\Delta\beta$ values that belong to each segment.

The inputs to the algorithm are the $\Delta\beta$ values $\Delta\beta = (\Delta\beta_1, .., \Delta\beta_i, .., \Delta\beta_N)$ to be segmented, the distance between adjacent CpGs $d = (d_1, ..., d_i, ..., d_N)$, the number of states $K^{(0)}$, the parameter $\omega$, the parameter $\theta$, and the distance normalization parameter $d_{Norm}$.

The meaning of the parameters $\theta$ and $\omega$ is the same we previously stated in Magi *et al.* (2011). The parameter $\omega$ modulates the proportionality between the white noise stochastic process ($\sigma_\epsilon^2$) and the means jump stochastic process ($\sigma_\mu^2$). For large values of $\omega$ (small $\sigma_\epsilon$ and large $\sigma_\mu$) the algorithm 'sees' the signal to have small white noise and takes as level shift also slight variations of the genomic profile. On the contrary when $\omega$ is small a large fraction of the total variance $\sigma^2$ is assigned to the $\sigma_\epsilon$ ($\sigma_\epsilon$ is large and $\sigma_\mu$ is small) and only sizeable variations of the signal are taken as level shift. The parameter $\theta$ corresponds to the baseline probability that a transition to a new mean level occurs at any position $i$ of the sequential process and is able to control only specificity and has weak effect on sensitivity. The parameter $K^{(0)}$ regulates the density of the $\mu_k$ in the range [-1,1]: the larger is $K^{(0)}$ and the larger is the $\mu_k$ density in the interval [-h,h]. The use of predefined $\mu_k$ values in the range [-1,1] does not affect the results of the segmentation procedure. This is due to the fact that the Viterbi algorithm is able to correctly identify all the mean shifts ($z_i$) also when the $\mu_k$ value does not perfectly match with the real value of the segments. In fact, when a state $m_i$ does not have its associated $\mu_k$, the Viterbi algorithm associates $m_i$ to the most lilkely $\mu_k$, allowing for the identification of all the mean shifts $z_i$. Finally, the distance normalization parameter $d_{Norm}$ modulates the ability of the segmentation algorithm to detect both small and highly isolated CpG of the genome and large and highly CpG-covered epi-genomic alterations. After segmentation, each segment is tested for differential methylation by using the Wilcoxon rank-sum test.

### Algorithm comparison

To test the ability of our segmentation algorithm to detect DMRs with different noise and degrees of methylation difference, we used the synthetic dataset generated in (Juhling *et al.*, 2016). In (Juhling *et al.*, 2016) the authors simulated DMRs on the human chromosome 10 (hg19) by exploiting the ENCODE chromatin state segmentation of the GM12878 cell line and using a Beta-Binomial approach to simulate both biological methylation and the sequencing step. By using a very complex recipe they simulated two different noise levels (low noise, background 1 and high noise, background 2) and four different degrees of methylation (with decreasing difference from class 1 to class 4) resulting in eight data sets with different levels of complexity, from easily (class 1 DMRs – background 1) to difficultly (class 4 DMR on background 2) distinguishable. For each of the height different level of complexity they simulated 20 samples, split in into two groups of 10 samples, and they introduced an equal number of hypermethylated DMRs (500) and hypomethylated DMRs (500).

To find the best parameter setting of our algorithm we applied it on the synthetic dataset of (Juhling *et al.*, 2016) by using $\omega = [0.1, 0.2, 0.3, 0.4, 0.5, 0.6, 0.7]$, $\theta = [10^-3, 10^-4, 10^-5, 10^-6, 10^-7]$ and $d_{Norm} = [10^4, 10^5, 10^6, 10^7]$. To use it on the analysis of multiple samples, for each CpG, we calculated the mean difference signal (MDS) with the following formula:

$$MDS_{CpG} = \frac{\sum_i \beta^i_{CpG}}{N} - \frac{\sum_j \beta^j_{CpG}}{M}. \quad (11)$$

where, for each CpG, $beta^i_{CpG}$ are the $\beta$ values of the N test samples and $beta^j_{CpG}$ are the $\beta$ values of the M test samples.

To evaluate the performance of our method we calculated precision as the ratio between the number of true positive (TP) DMRs and the total number of called DMRs and recall as the ratio between the number of TP and the total number of simulated DMRs: a detected segment is considered a TP if there is at least a 50% overlap with the simulated DMRs. The results reported in Supplemental Figure 1-6 show that for low complexity datasets (low noise and high methylation differences) our method obtains the best results with $\omega = [0.2, 0.3]$ and $\eta = [10^-4, 10^-3]$, while for high complexity dataset (high noise and low methylation differences) the best F-score is obtained with $\omega = [0.5, 0.6]$ and $\eta = [10^-7, 10^-6]$. Concerning the $d_{Norm}$ parameter, it has no effect on the global performance

As a further step we compared the performance of our method with those of two previously published algorithms: Metilene (Juhling *et al.*, 2016) and BSmooth (Hansen *et al.*, 2012). We downloaded Metilene version 0.2-8 from `https://www.bioinf.uni-leipzig.de/Software/metilene/Downloads/` and the bsseq R package (that contains the BSmooth algorithm) from Bioconductor and we ran them with default parameters settings. The results of Supplemental Figure 8 demonstrate that our segmentation method outperforms the other two state-of-the-art methods for all the class of data complexity (high and low noise) especially in the analysis of a small number of samples (Supplemental Figure 8.e-f).

### Visual inspection

Visual inspection of aligned reads is a fundamental step in the identification of genomic variants since the advent of SGS technologies. The discovery of putative variants is performed by automated computational pipelines, but visual review remains critical to the verification and interpretation of these variants (Robinson *et al.*, 2017), especially for SVs with long read data that are inferred by using a very complex combination of alignment signatures (gapped alignment and split read alignment or read-depth).

Although NanomonSV combines gapped alignment and split read alignment signatures, small deletions and insertions ($< 5$ kb) are generally inferred by using gapped alignment signatures, while inversions, duplication, translocations and large insertions and deletions (larger than 5 kb) are detected by exploiting split read signature.

For these reasons, small deletions and insertions were visually validated by using the Integrative Genomics Viewer (IGV version 2.9.4) (Robinson *et al.*, 2011) while duplications, inversions and large insertions and deletions by using Samplot version 1.3.0 (Belyeu *et al.*, 2021).

IGV was one of the first tools to provide SGS data visualization and recently has been updated with a suite of specialized features to support third-generation long-read sequencing technologies. Samplot is a command line tool that allows to produce high-quality images that highlight any split-alignment and depth signals that support SV.
The results of visual inspection of the SVs identified by NanomonSV in the three AML pairs are reported in Supplemental File 4.

Table 1: AML samples characteristics.

| Sample | Sex | Age | FAB | Karyotype | Molecular profile | 1st line treatment (induction, maintenance) |
|---|---|---|---|---|---|---|
| AML2 | F | 20 | M2 | XX,46 | STAG2 | Idarubicin + HiDAC, AraC |
| UD5 | M | 67 | myelodysplastic changes | XY,46 | NPM1 | FLAI, Idarubicin + AraC, HiDAC + autoHSCT |
| UD10 | F | 22 | M4 | inv(16) | NA | FLAI, Idarubicin + AraC, HiDAC |

The table reports the main features at diagnosis of the three AML patients: Sex, Age, French-American-British (FAB) classification, the Karyotype, the Molecular profile and the 1st line treatment. Molecular profile was obtained through Illumina targeted sequencing of bone marrow aspirate performed as per clinical routine. NA, not available; HiDAC, high-dose cytarabine; AraC, standard-dose cytarabine; FLAI, fludarabine, cytarabine, idarubicin; autoSCT, autologous hematopoietic stem cell transplantation.

Table 2: CpG groups inferred by Nanopolish.

| Sample | CpG Groups | 1 CpG site | 2 CpG sites | 3 CpG sites | 4 CpG sites | ≥5 CpG sites | Total CpG sites |
|---|---|---|---|---|---|---|---|
| UD5T | 22687510 | 19191168 (84.5%) | 2461372 (10.9%) | 635351 (2.8%) | 217004 (1%) | 182615 (0.8%) | 28181699 |
| UD5R | 22689245 | 19192734 (84.5%) | 2461413 (10.9%) | 635404 (2.8%) | 217043 (1%) | 182651 (0.8%) | 28183843 |
| UD10T | 22578261 | 19093348 (84.5%) | 2452640 (10.9%) | 633692 (2.8%) | 216224 (1%) | 182357 (0.8%) | 28056760 |
| UD10R | 22574289 | 19089697 (84.5%) | 2452425 (10.9%) | 633637 (2.8%) | 216184 (1%) | 182346 (0.8%) | 28052309 |
| AML2T | 22531536 | 19058637 (84.5%) | 2445007 (10.9%) | 631044 (2.8%) | 215123 (1%) | 181725 (0.8%) | 27990455 |
| AML2R | 22529573 | 19057120 (84.5%) | 2444698 (10.9%) | 630949 (2.8%) | 215116 (1%) | 181690 (0.8%) | 27987801 |

The table reports the total number CpG groups inferred by Nanopolish in each sample (CpG Groups). Columns from 3 to 7 report the number CpG groups containing 1 (1 CpG site), 2 (2 CpG sites), 3 (3 CpG sites), 4 (4 CpG sites) and more than five (≥5 CpG sites) CpG sites. Last column reports the cumulative number of CpG sites in all CpG groups.

Table 3: DMRs annotation statistics

| Sample | # DMRs | # DHS | # TFBS | Enhancers | # Genes | # CGIs | # NoCGIs | # Hyper-CGIs | # Hyper-NoCGIs | # Hypo-CGIs | Hypo-NoCGIs |
|---|---|---|---|---|---|---|---|---|---|---|---|
| UD5 | 1269 | 1206 (95%) | 1052 (83%) | 199 (16%) | 919 (72%) | 325 (26%) | 325 (74%) | 944 (96%) | 312 (4%) | 13 (54%) | 510 (46%) |
| UD10 | 1149 | 1111 (97%) | 1017 (89%) | 118 (10%) | 822 (72%) | 744 (65%) | 744 (35%) | 405 (97%) | 718 (3%) | 26 (63%) | 254 (37%) |
| AML2 | 1314 | 1257 (96%) | 1155 (88%) | 207 (16%) | 1014 (77%) | 596 (45%) | 596 (55%) | 718 (98%) | 586 (2%) | 10 (86%) | 620 (14%) |

The table reports statistics about DMRs annotation. For each sample (UD5, UD10 and AML2) are reported the total number of DMRs (# DMRs), the number of DMRs overlapping DHS (# DHS), TFBS (# TFBS), annotated genes (# Genes), CpG islands (# CGIs), and sparse CpGs (# NoCGIs, DMRs that do not overlap any CGIs). The last four columns report the number of number of hyper- and hypo-methylated DMRs overlapping a not overlapping CGIs (# Hyper-CGIs, # Hyper-NoCGIs, # Hypo-CGIs, # Hypo-NoCGIs). Percentages in brackets show proportion with respect to total DMRs. For the last four columns percentages report proportion with respect to total hyper- (# Hyper-CGIs, # Hyper-NoCGIs ) and hypo-methylated (# Hypo-CGIs, # Hypo-NoCGIs) DMRs.

Table 4: Differentially expressed Genes in the three AML patients.

| Sample | DEGs | Over-expressed Genes | Under-Expressed Genes |
|---|---|---|---|
| UD5 | 4,677 | 2,044 | 1,953 |
| UD10 | 3,997 | 2,890 | 1,787 |
| AML2 | 1,759 | 495 | 1,264 |

The table reports the total number of differentially expressed genes (DEGs), over-expressed and under-expressed genes identified by DESeq2 (adjusted p-value$< 0.05$ and absolute $log_2FC > 0.5$) on the three AML samples.

Table 5: Differentially methylated and Differentially expressed Genes in the three AML patients.

| Sample | Hyper-Over | Hypo-Over | Hyper-Under | Hypo-Under |
|---|---|---|---|---|
| UD5 | 97 (5%) | 40 (2%) | 100 (5%) | 39 (2%) |
| UD10 | 33 (1%) | 22 (1%) | 51 (3%) | 17 (1%) |
| AML2 | 19 (4%) | 5 (1%) | 75 (6%) | 5 (0%) |

The table reports the total number of Differentially methylated and Differentially expressed Genes (DM-DEGs) for the three AML samples. Statistics are reported for the four methylation-expression combination: hyper-methylation and over-expression (Hyper-Over), hypo-methylation and over-expression (Hypo-Over), hyper-methylation and under-expression (Hyper-Under), hypo-methylation and under-expression (Hypo-Under). For each category, percentages in brackets report the proportions of DM-DEGs with respect to total number of over- or under-expressed genes.

Table 6: Relapse-specific SVs detected by NanomonSV in the three AML pairs.

| Chr | Start | End | Size | Type | RS | TS | Sample | Gene | Visual Inspection | Tool | Figure |
|---|---|---|---|---|---|---|---|---|---|---|---|
| 3 | 116259740 | 116486402 | 235795 | Del | 7/19 (0.36) | 0/77 | UD5 | TUSC7 (All) | Relapse-specific | Samplot | 1 |
| 3 | 76182548 | 76182814 | 266 | Del | 4/32 (0.12) | 0/28 | UD5 | Yes ROBO2 (Intron) | Relapse-specific | IGV | 2 |
| 3 | 116604969 | 116631956 | 26987 | Dup | 9/63 (0.14) | 0/72 | UD5 | | Relapse-specific | Samplot | 3 |
| 7 | 115992315 | 115994264 | 1949 | Del | 9/40 (0.22) | 0/38 | UD5 | CAV2 (Exon) | Relapse-specific | IGV | 4 |
| 16 | 50350527 | 50352587 | 2060 | Del | 9/34 (0.26) | 0/26 | UD5 | ADCY7 (Exon) | Relapse-specific | IGV | 5 |
| 6 | 32454684 | 32478662 | 23978 | Del | 5/25 (0.2) | 0/36 | UD10 | | Not Relapse-specific | IGV | 6 |
| 13 | 35684164 | 35684406 | 242 | Del | 4/25 (0.16) | 0/28 | UD10 | NBEA (Intron) | Relapse-specific | IGV | 7 |
| 17 | 20707099 | 20707205 | 106 | Del | 3/26 (0.11) | 0/24 | UD10 | | Relapse-specific | IGV | 8 |
| 19 | 43718061 | 43718371 | 310 | Del | 4/13 (0.3) | 0/10 | UD10 | PSG9 (Intron) | Relapse-specific | IGV | 9 |
| 4 | 140244927 | 143675814 | 3430887 | Del | 7/34 (0.2) | 0/51 | AML2 | SETD7, MGST2, MAML3, SCOC, ELMOD2, UCP1, ZNF330, IL15, INPP4B (All) | Relapse-specific | Samplot | 10 |
| 14 | 106215508 | 106215781 | 273 | Del | 3/21 (0.14) | 0/21 | AML2 | IGHM (Intron) | Relapse-specific | IGV | 11 |
| 19 | 23647827 | 23648154 | 327 | Del | 4/19 (0.21) | 0/19 | AML2 | | Relapse-specific | IGV | 12 |
| 19 | 23655167 | 23655537 | 370 | Del | 3/15 (0.2) | 0/16 | AML2 | | Relapse-specific | IGV | 13 |

Columns report the genomic coordinates (Chr, Start and End, with respect to hg19), the size (in bp), the type of SV (Del, Dup), the number of signatures supporting the SV and the total number of reads with reference allele in the relapsed sample (RS, signature number are separated by /, between brackets is reported cellular fraction), the number of signatures supporting the SV and the total number of reads with reference allele in the tumor sample (TS, signature number are separated by /), the sample name, the results obtained by NanomonSV in the identification of the SV ,the result of visual inspection (Relapse-specific if SV signatures are present only in relapse sample, Not Relapse-specific if SV signatures are present in both relapsed and tumor samples), the visualization tool used for visual inspection (Tool, Samplot or IGV) and the figure number in which are reported the results of visual inspection in supplemental file 4.

Table 7: Variants identified by WES in the three AML samples.

| Sample | Disease Phase | Gene | Chr | Start | End | Ref | Alt | Variant Type | $T_{VAF}$ | $R_{VAF}$ |
|---|---|---|---|---|---|---|---|---|---|---|
| UD5 | T | MYPOP | 19 | 46394248 | 46394248 | C | T | nonsynonymous SNV | 12.07 | - |
| UD5 | R/T | BRINP2 | 1 | 177250213 | 177250213 | G | A | nonsynonymous SNV | 44.63 | 45.38 |
| UD5 | R/T | ZC3HC1 | 7 | 129679315 | 129679315 | C | A | nonsynonymous SNV | 38.85 | 45.40 |
| UD5 | R/T | FILIP1L | 3 | 99648839 | 99648839 | T | C | nonsynonymous SNV | 46.46 | 49.00 |
| UD5 | R/T | VN1R2 | 19 | 53762305 | 53762305 | T | G | nonsynonymous SNV | 43.14 | 42.52 |
| UD5 | R/T | CEBPA | 19 | 33792821 | 33792821 | - | C | frameshift insertion | 11.11 | 45.83 |
| UD5 | R/T | DSPP | 4 | 88536863 | 88536880 | AATAGTAGTGACAGCAGC | - | nonframeshift deletion | 70.49 | 89.47 |
| UD5 | R/T | NMNAT1 | 1 | 10035794 | 10035794 | - | T | frameshift insertion | 39.07 | 43.31 |
| UD5 | R/T | NPM1 | 5 | 170837543 | 170837543 | - | TCTG | frameshift insertion | 47.06 | 45.45 |
| UD5 | R/T | PRR21 | 2 | 240982109 | 240982136 | GAAGGGCCGTGGGTGAAGAGGCATGGAT | - | frameshift deletion | 43.48 | 42.86 |
| UD5 | R | SCIN | 7 | 12691477 | 12691477 | G | A | stopgain | - | 10.00 |
| UD5 | R | SLC44A1 | 9 | 108118642 | 108118642 | G | A | nonsynonymous SNV | - | 11.11 |
| UD5 | R | KRT23 | 17 | 39081765 | 39081765 | T | A | nonsynonymous SNV | - | 20.00 |
| UD5 | R | SLC2A3 | 12 | 8084037 | 8084037 | C | A | nonsynonymous SNV | - | 21.25 |
| UD5 | R | ZNF34 | 8 | 145998937 | 145998937 | T | C | nonsynonymous SNV | - | 25.00 |
| UD5 | R | GPR123 | 10 | 134942360 | 134942360 | G | C | nonsynonymous SNV | - | 30.60 |
| UD5 | R | ENDOD1 | 11 | 94861922 | 94861922 | C | A | nonsynonymous SNV | - | 30.88 |
| UD5 | R | CTNNA2 | 2 | 80136765 | 80136765 | C | T | nonsynonymous SNV | - | 44.83 |
| UD5 | R | OR8B8 | 11 | 124310617 | 124310617 | C | T | nonsynonymous SNV | - | 18.62 |
| UD5 | R | TTN | 2 | 179584137 | 179584137 | C | T | nonsynonymous SNV | - | 15.15 |
| UD5 | R | FLT3 | 13 | 28608268 | 28608268 | - | TCTCTGAAATCAACGTAGAAGTACTCATTATCTGAGGAGCCGGTCACCTGTACCATCTGA | nonframeshift insertion | - | 13.30 |
| UD10 | R/T | NRAS | 1 | 115256530 | 115256530 | G | T | nonsynonymous SNV | 41.90 | 40.00 |
| UD10 | R/T | YLPM1 | 14 | 75248804 | 75248804 | T | A | nonsynonymous SNV | 40.00 | 45.10 |
| UD10 | R/T | KRT8 | 12 | 53298675 | 53298675 | A | C | nonsynonymous SNV | 13.04 | 11.76 |
| UD10 | R/T | CTNNA3 | 10 | 68526128 | 68526128 | G | A | nonsynonymous SNV | 42.00 | 44.78 |
| UD10 | R/T | ISLR | 15 | 74468241 | 74468241 | G | A | nonsynonymous SNV | 42.96 | 41.03 |
| UD10 | R/T | SYT10 | 12 | 33559867 | 33559867 | G | A | stopgain | 48.33 | 46.67 |
| UD10 | R | TMEM132D | 12 | 129566312 | 129566312 | T | G | nonsynonymous SNV | 2.38 | 10.00 |
| UD10 | R | IVL | 1 | 152883532 | 152883532 | T | G | nonsynonymous SNV | 3.85 | 19.51 |
| UD10 | R | KDR | 4 | 55970945 | 55970945 | A | G | nonsynonymous SNV | - | 30.97 |
| UD10 | R | TRPC4 | 13 | 38320085 | 38320085 | G | A | nonsynonymous SNV | - | 34.90 |
| UD10 | R | DMKN | 19 | 36002421 | 36002421 | - | CTGCTGCTG | nonframeshift insertion | - | 57.14 |
| UD10 | R | DSPP | 4 | 88537205 | 88537213 | GACAGCAGT | - | nonframeshift deletion | - | 78.95 |
| AML2 | T | GOLGA6L9 | chr15 | 82434465 | 82434465 | C | T | stopgain | 11.50 | - |
| AML2 | T | PDPR | chr16 | 70144498 | 70144498 | A | T | nonsynonymous SNV | 35.70 | - |
| AML2 | R/T | TCERG1L | chr10 | 131260421 | 131260421 | C | A | nonsynonymous SNV | 59.00 | 55.40 |
| AML2 | R/T | NUAK1 | chr12 | 106066930 | 106066930 | G | T | nonsynonymous SNV | 48.30 | 44.70 |
| AML2 | R/T | PCDH8 | chr13 | 52847542 | 52847542 | G | A | nonsynonymous SNV | 47.40 | 49.50 |
| AML2 | R/T | KIAA1755 | chr20 | 38217300 | 38217300 | G | A | nonsynonymous SNV | 45.20 | 45.80 |
| AML2 | R/T | TBXAS1 | chr7 | 139953419 | 139953419 | C | T | nonsynonymous SNV | 44.90 | 48.50 |
| AML2 | R/T | RNF20 | chr9 | 101554106 | 101554106 | G | A | splice$_{s}ite$ | 46.60 | 44.20 |
| AML2 | R/T | STAG2 | chrX | 124045321 | 124045321 | T | A | stopgain | 48.20 | 45.70 |
| AML2 | R | POTEI | chr2 | 130463831 | 130463831 | G | C | nonsynonymous SNV | - | 47.90 |
| AML2 | R | ANAPC4 | chr4 | 25418249 | 25418259 | GTAATAAGCCT | - | frameshift deletion | - | 32.80 |

Columns report the sample name (Sample), the Disease Phase in which the variant has been detected (T for primary tumor, R for relapse, R/T for common to tumor and relapse), the gene affected by the variant (Gene), the genomic coordinates (Chr, Start and End, with respect to hg19), the reference and alternative alleles that describe the variant (Ref and Alt), the functional effect (Variant Type) and the variant allele fraction in primary tumor ($T_{VAF}$) and in relapsed samples ($R_{VAF}$).

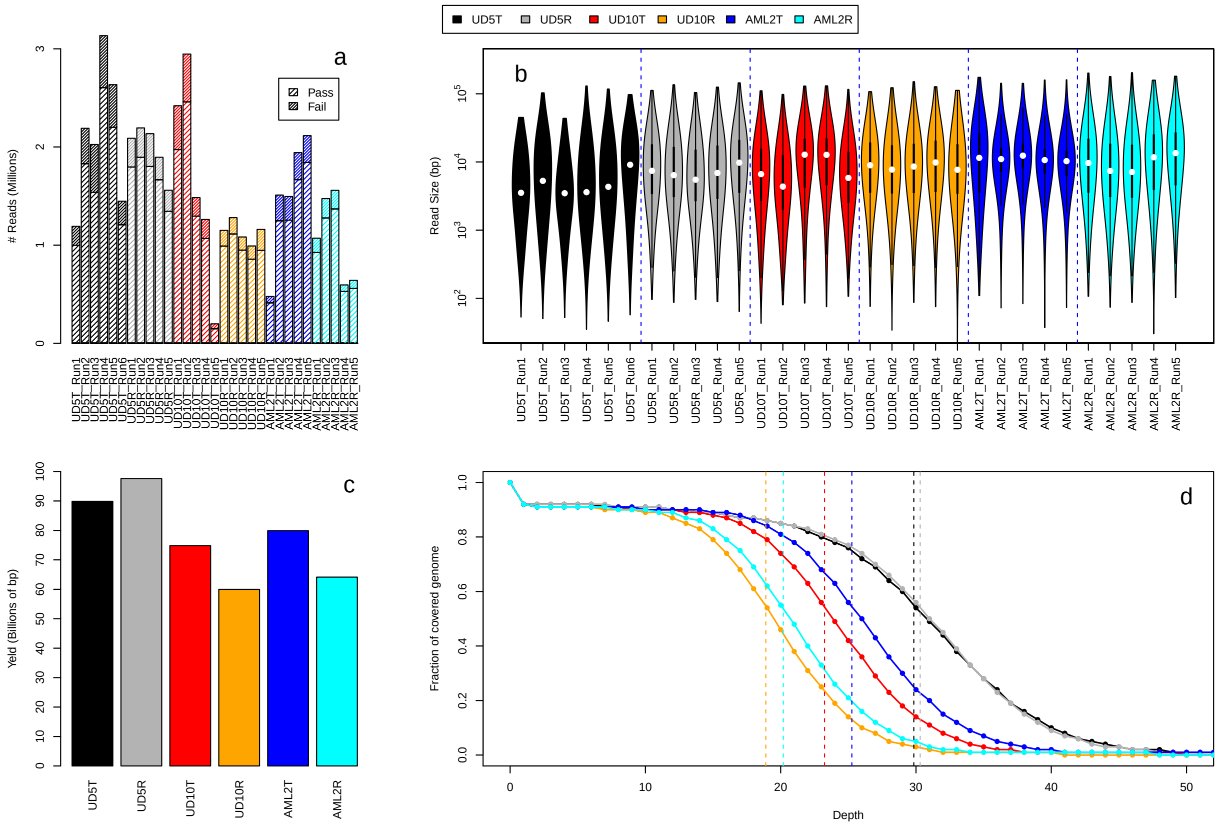


Figure 1: Sequencing Statistics and Structural variants. Panel (a) shows the total number of reads (Pass and Fail) generated by the 31 sequencing run performed on the six AML samples. Each sequencing run generated between 250000 and 3.2 millions total reads, with a fraction of low-quality reads (fail) around 15-25% that were discarded by the base-caller. The violin plots of panel (b) report the sequence size distribution of pass reads for each sequencing run. Pass read size distributions were consistent across all the runs with a mean read length that ranges between a minimum of 5857 bp (UD5T_Run3) to a maximum of 17584 bp (AML2R_Run5) For each sample, panel (c) shows the total sequencing throughput of pass reads. Panel (d) reports the fraction of genome as a function of sequencing coverage obtained by minimap2. Vertical dotted lines indicate the average coverage for each sequenced sample: 30x (UD5T), 30x (UD5R), 24x (UD10T), 19x (UD10R), 25x (AML2T) and 20x (AML2R).

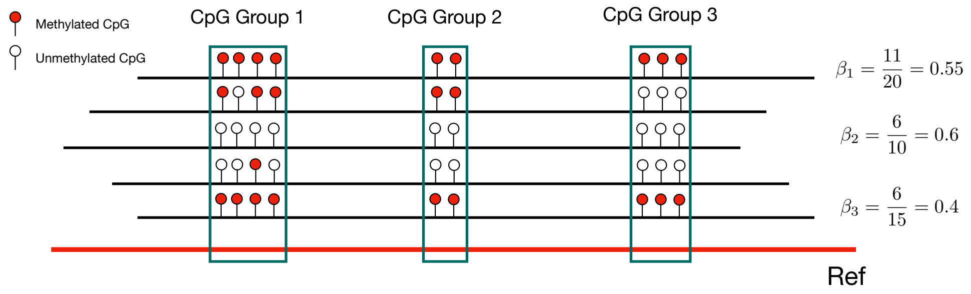


Figure 2: $\beta$ values calculation. Nanopolish is based on a hidden Markov model that assigns a log-likelihood ratio to a group of nearby CpG sites that share the same methylation level in each site within the group. To estimate the methylation level ($\beta$), for each CpG group identified by Nanopolish, we calculated the ratio between the total number of CpG sites predicted as methylated divided by the total number of CpG sites in the group. In figure are reported $\beta$ calculation of three CpG groups inferred by Nanopolish. Red line indicates the reference genome, black lines indicate the reads, red (white) dots indicate methylated (unmethylated) CpGs inferred by Nanopolish.

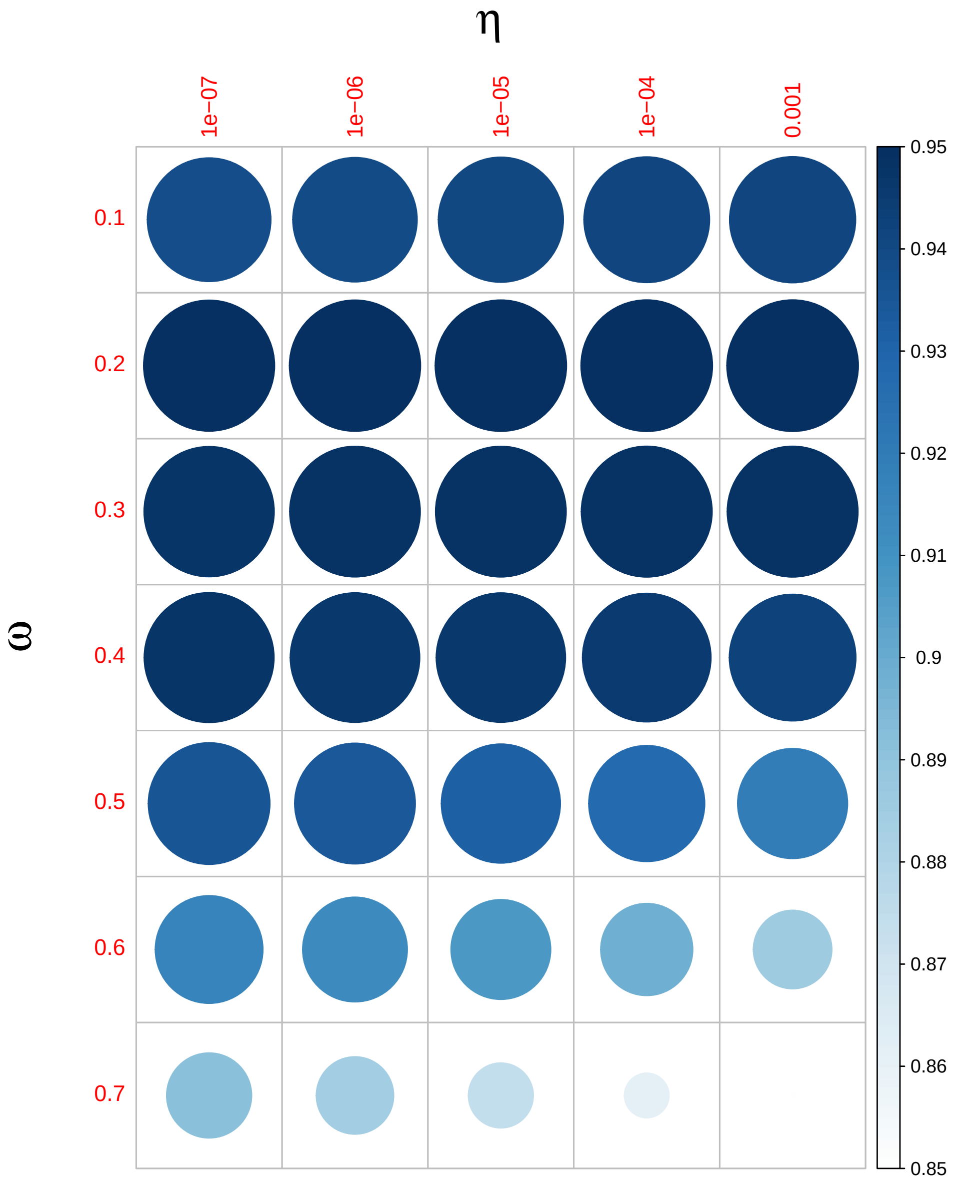


Figure 3: PoreMeth and parameter settings on synthetic data. The dotplot shows the F-score (harmonic mean of the precision and recall) obtained by PoreMeth on the analysis of synthetic data with background 1, for different combinations of $\omega$ and $\theta$.

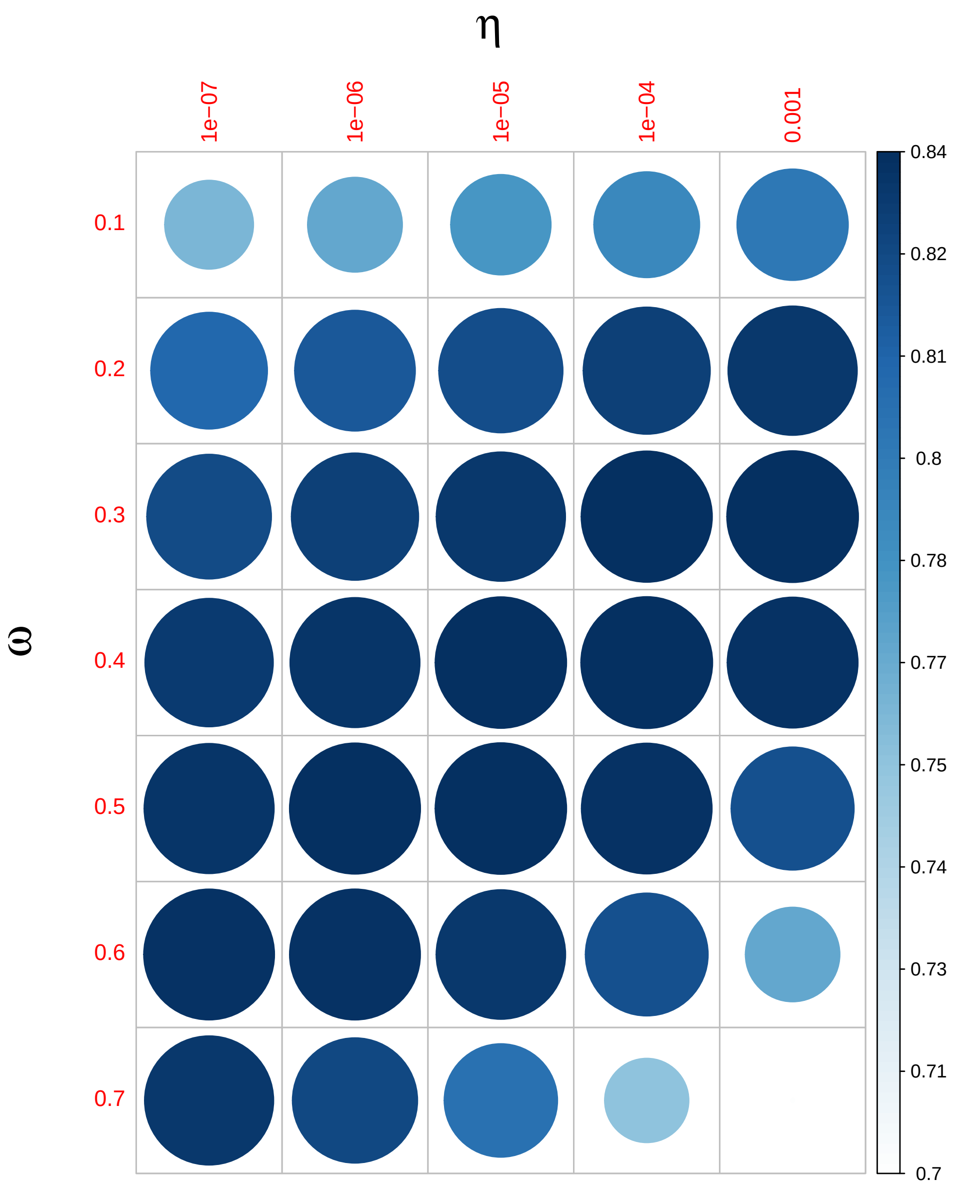


Figure 4: PoreMeth and parameter settings on synthetic data. The dotplot shows the F-score (harmonic mean of the precision and recall) obtained by PoreMeth on the analysis of synthetic data with background 2, for different combinations of $\omega$ and $\theta$.

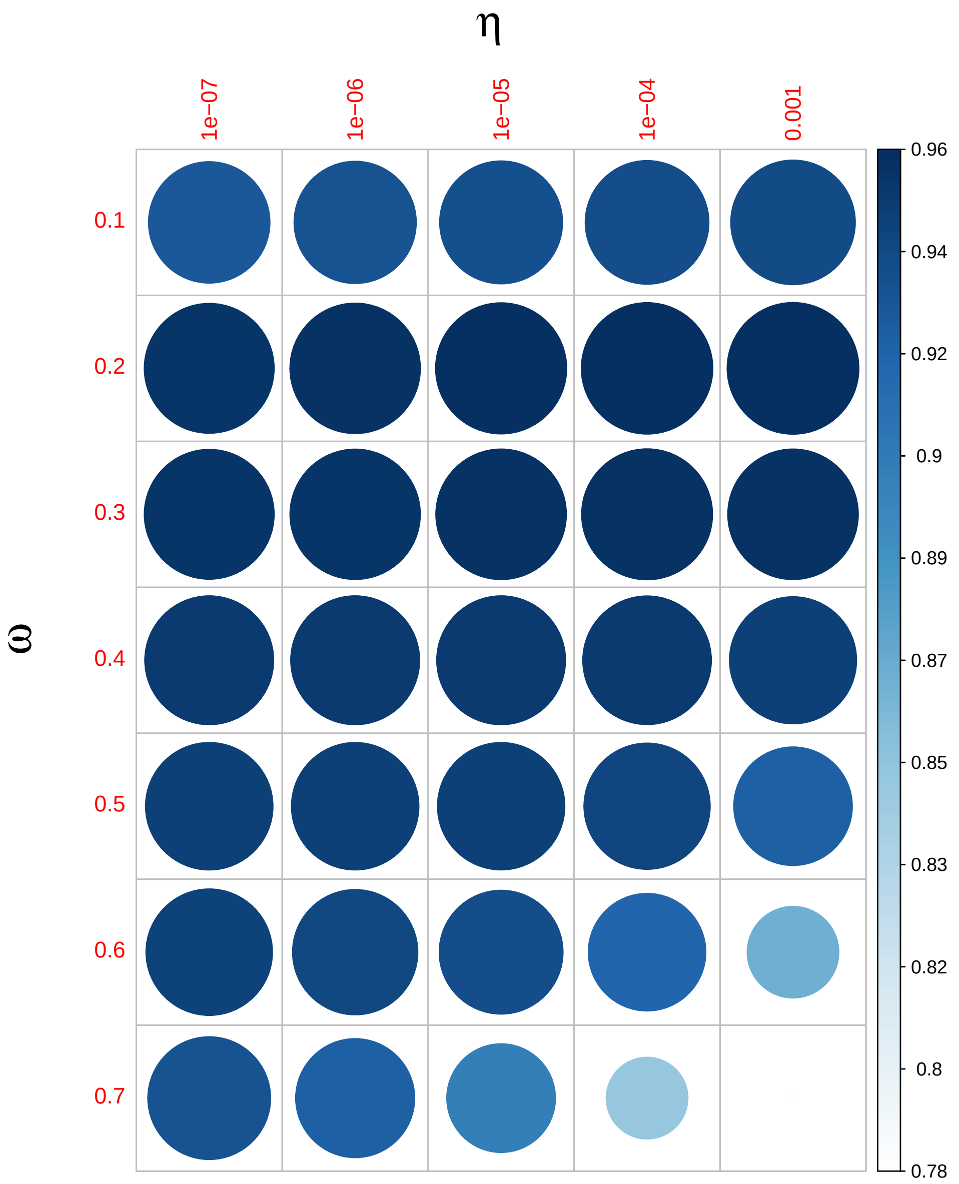


Figure 5: PoreMeth and parameter settings on synthetic data. The dotplot shows the F-score (harmonic mean of the precision and recall) obtained by PoreMeth on the analysis of synthetic data with degrees of methylation of class 1, for different combinations of $\omega$ and $\theta$.

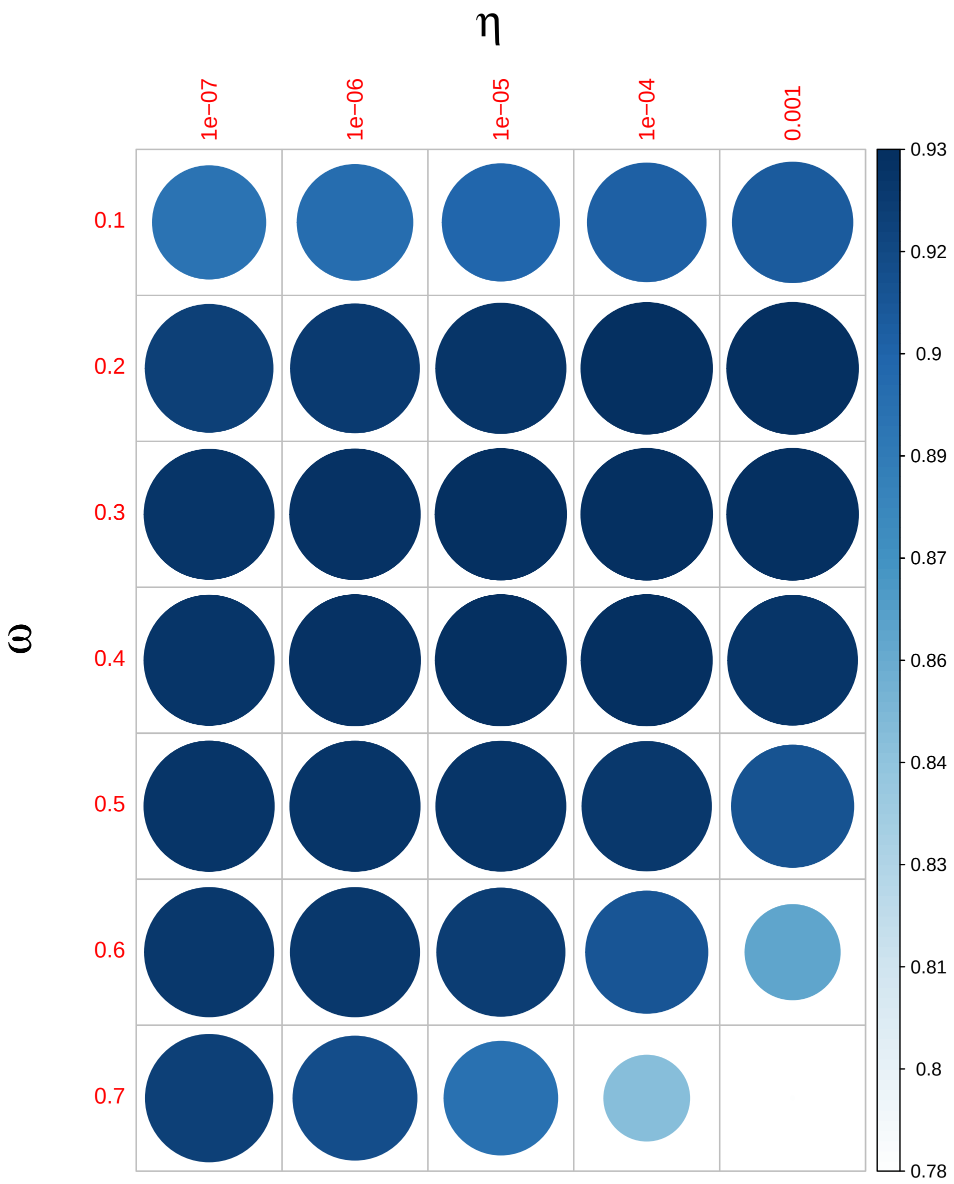


Figure 6: PoreMeth and parameter settings on synthetic data. The dotplot shows the F-score (harmonic mean of the precision and recall) obtained by PoreMeth on the analysis of synthetic data with degrees of methylation of class 2, for different combinations of $\omega$ and $\theta$.

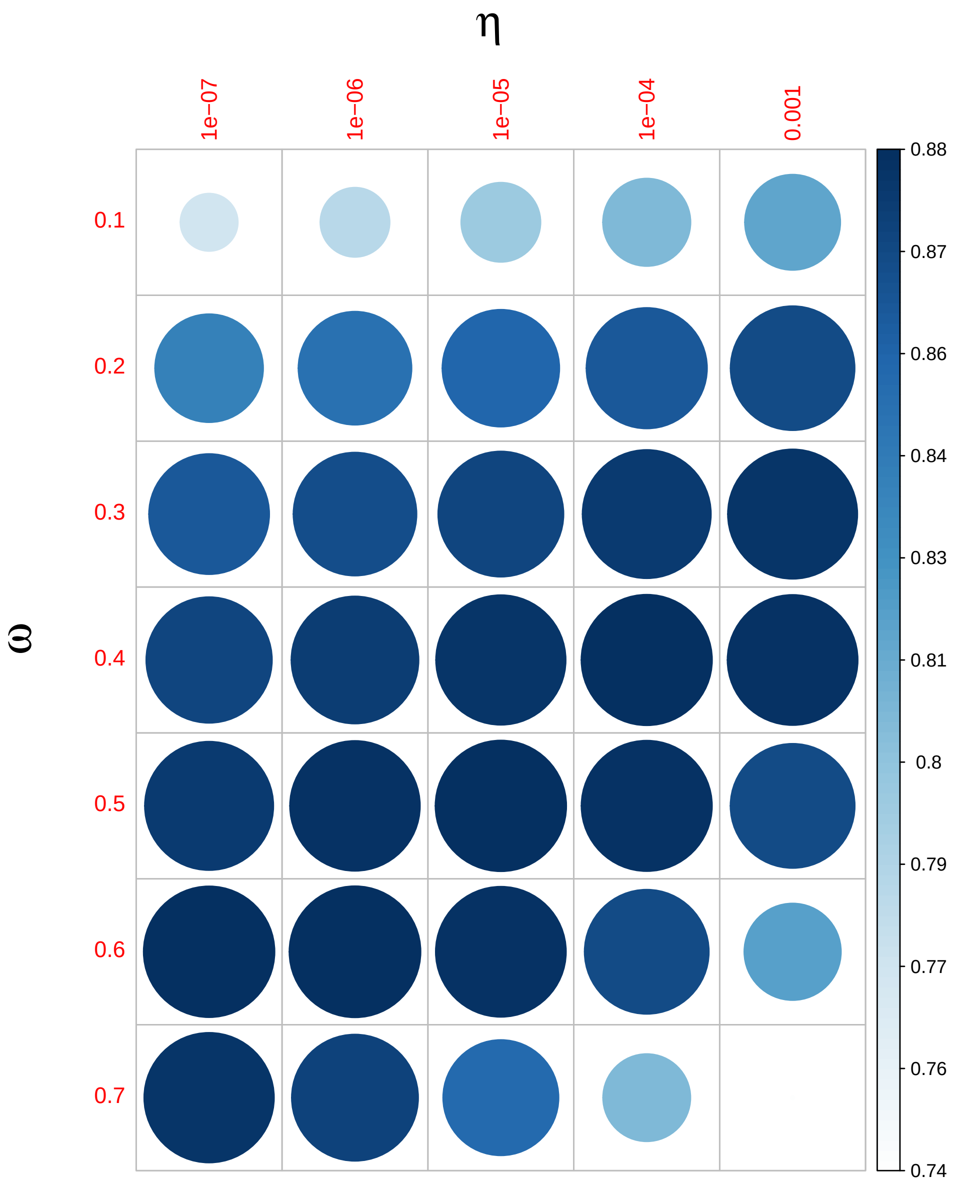


Figure 7: PoreMeth and parameter settings on synthetic data. The dotplot shows the F-score (harmonic mean of the precision and recall) obtained by PoreMeth on the analysis of synthetic data with degrees of methylation of class 3, for different combinations of $\omega$ and $\theta$.

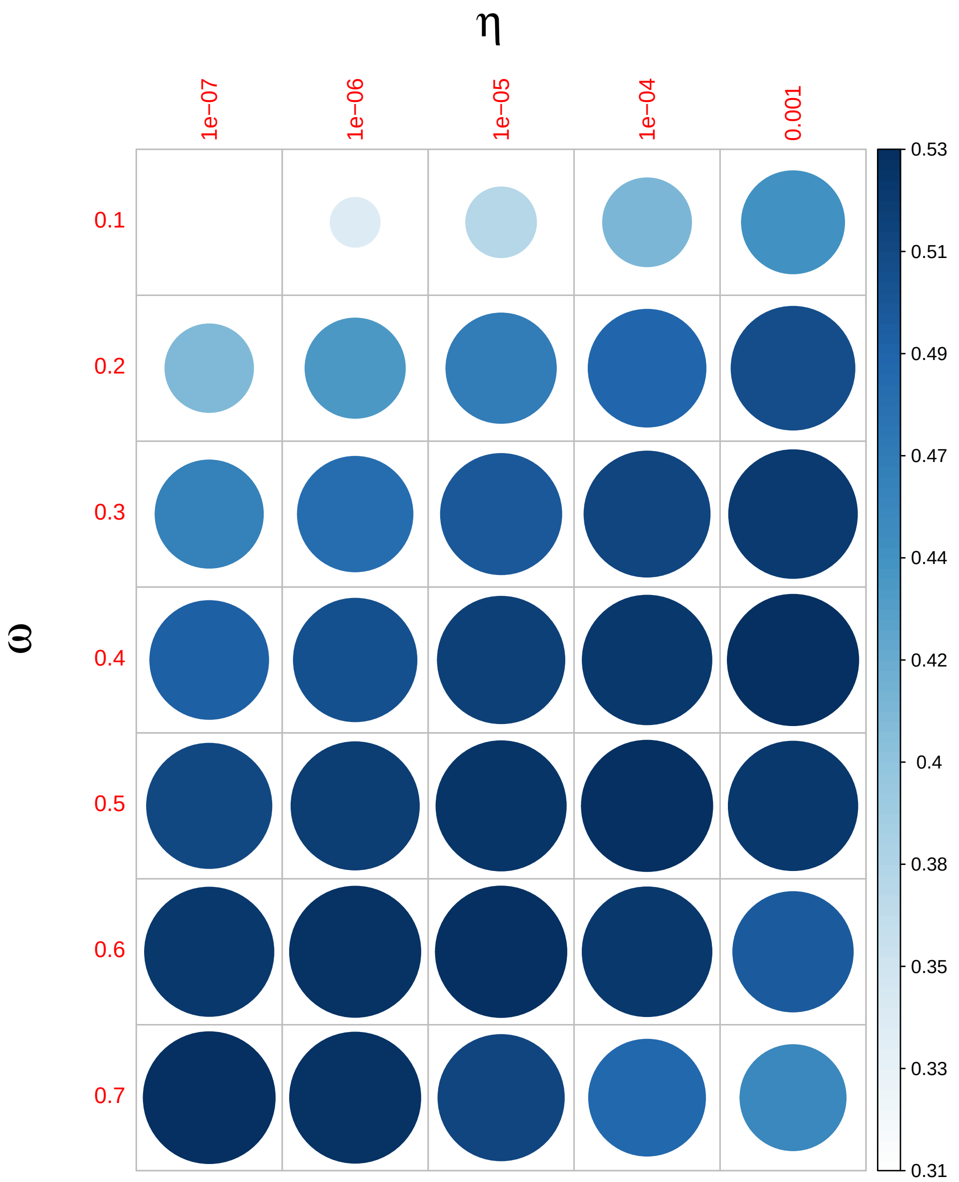


Figure 8: PoreMeth and parameter settings on synthetic data. The dotplot shows the F-score (harmonic mean of the precision and recall) obtained by PoreMeth on the analysis of synthetic data with degrees of methylation of class 4, for different combinations of $\omega$ and $\theta$.

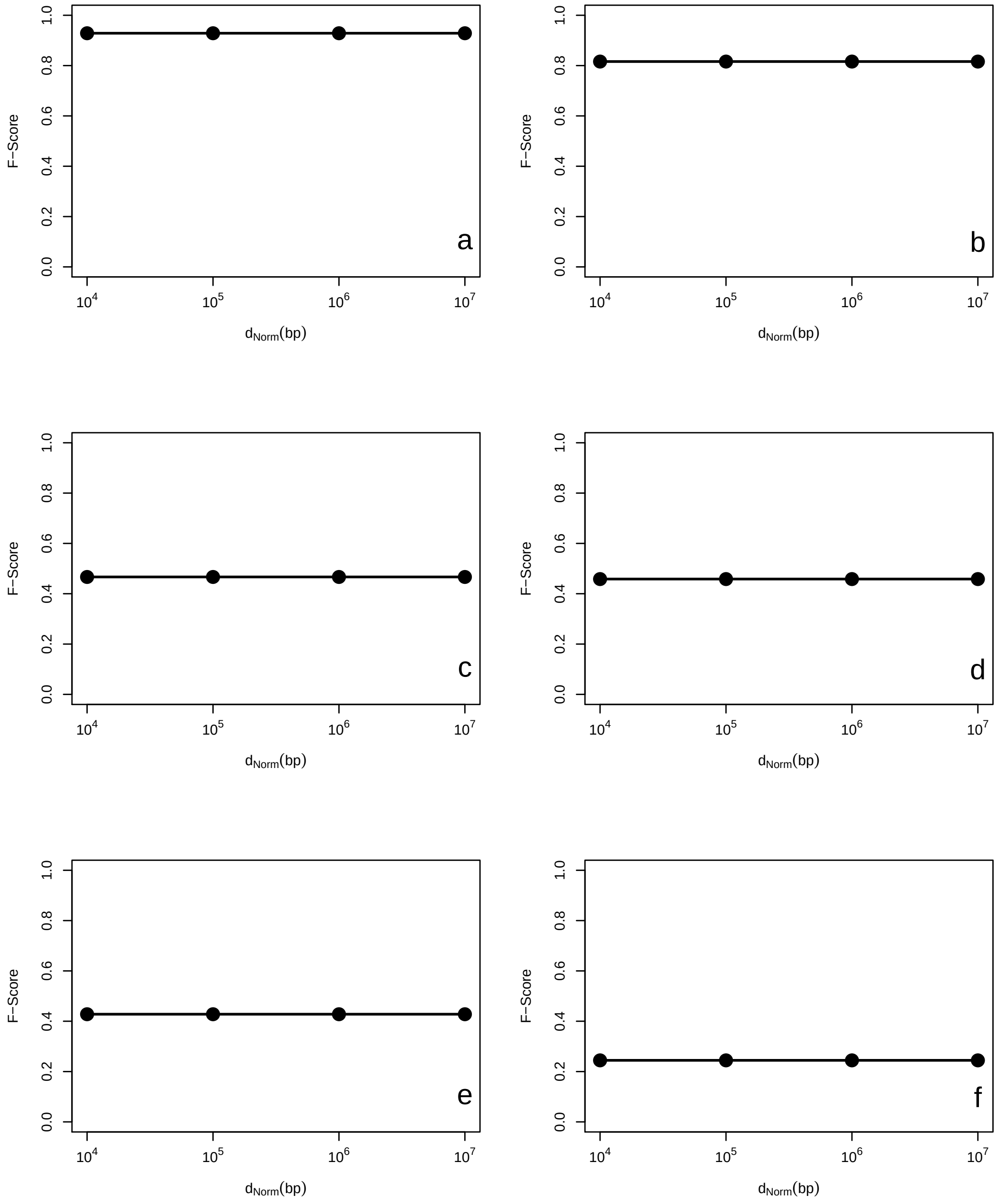


Figure 9: PoreMeth and parameter settings on synthetic data. The plot shows the F-score (harmonic mean of the precision and recall) obtained by PoreMeth on the analysis of synthetic data with background 1-2 (panels a-b) and degrees of methylation of class 1-4 (panels c-f), for different values of $d_{Norm}$.

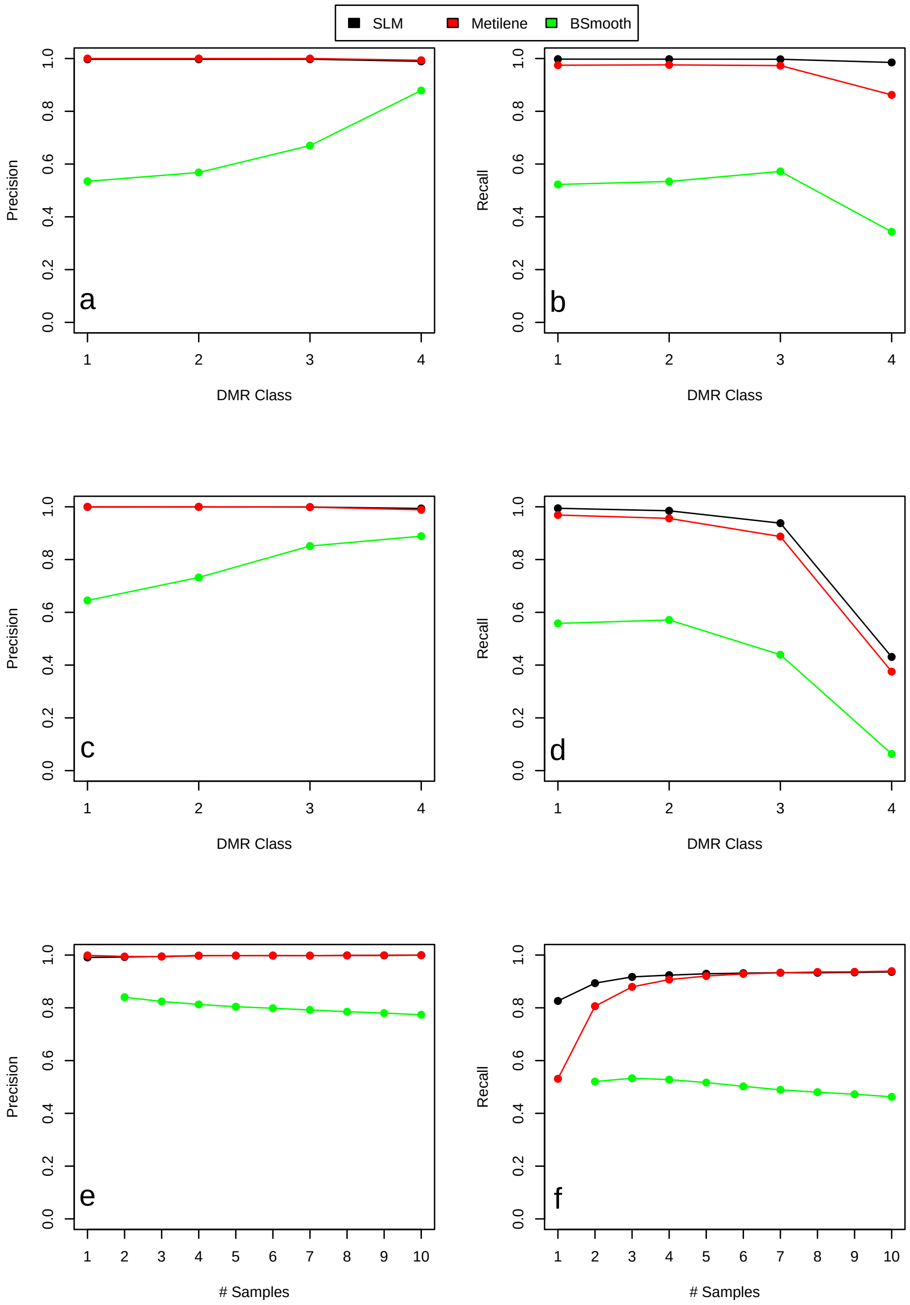


Figure 10: Comparison between PoreMeth, Metilene and BSmooth.The plot shows precision and recall obtained by the three tools as a function of differential methylation class (a-d) and as a function of analyzed samples (e-f). Panels a-b report the results for background 1 (low noise), while panels c-d for background 2 (high noise).

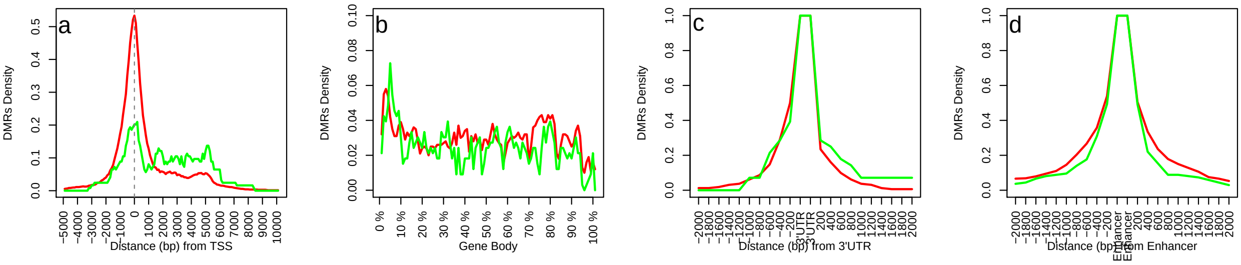


Figure 11: DMRs distribution across genomic. The four panels report the distribution of DMRs at 5' regulatory region (a), at gene-body (b), at 3'UTR (c) and at enhancers (d). Red lines for hyper-methylation and green lines for hypo-methylation.

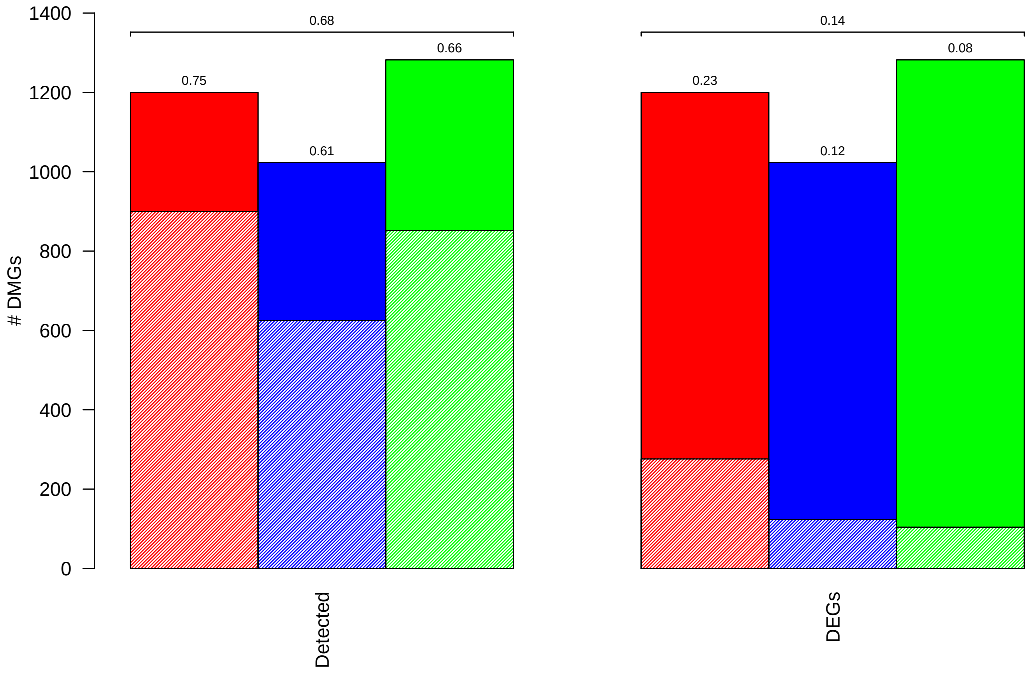


Figure 12: Differentially-methylated genes (DMGs) and RNASeq results on the three AML pairs. Left barplots report the total number of DMGs (solid bars) and the numbers of DMGs with detectable RNASeq signals (textured bars). Right barplots report the total number of DMGs (solid bars) and the number of DMGs that are also differentially expressed (textured bars, adjusted p-value$< 0.05$ and absolute log2FC $> 0.5$) in the three AML pairs. Numbers above bars show the percentage of DMGs with detectable RNASeq signals (left) and DM-DEGs (right).

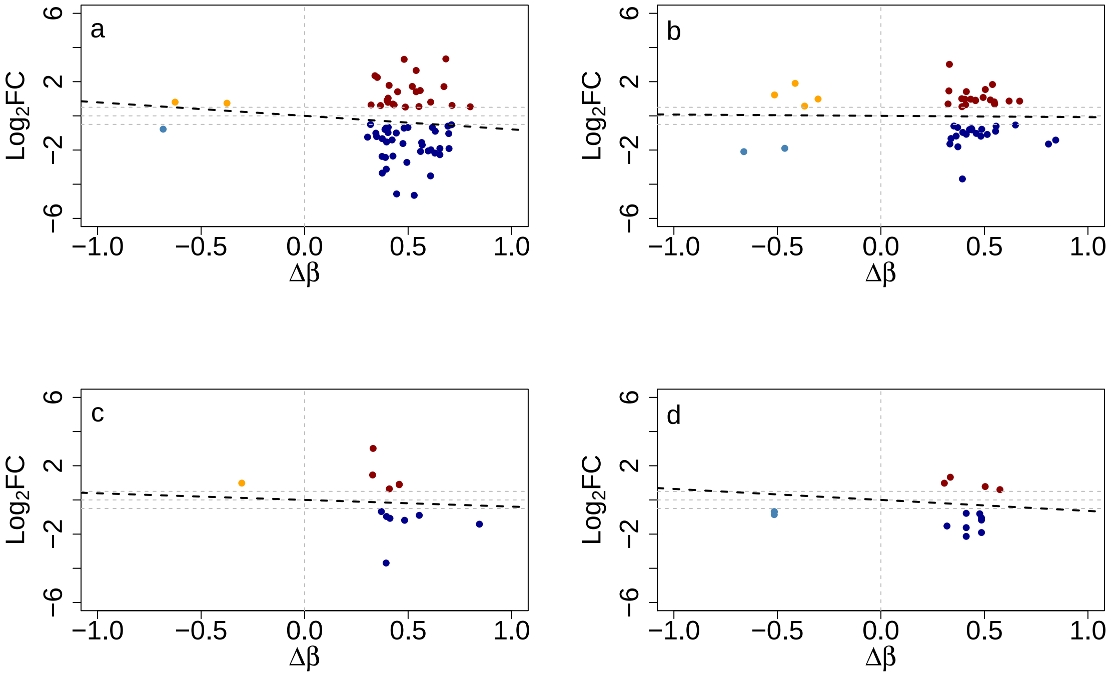


Figure 13: Correlation between $Log_2FC$ and $\Delta\beta$ values in genomic elements that overlap CpG islands. Panels show the correlation between $Log_2FC$ and $\Delta\beta$ values of DEG with differentially methylated regulatory element (a), gene body (b), 3'UTR (c) and enhancer (d). All the DMRs taken into consideration in this analysis overlap with a CpG island. Dotted line represent the fitted linear regression curve.

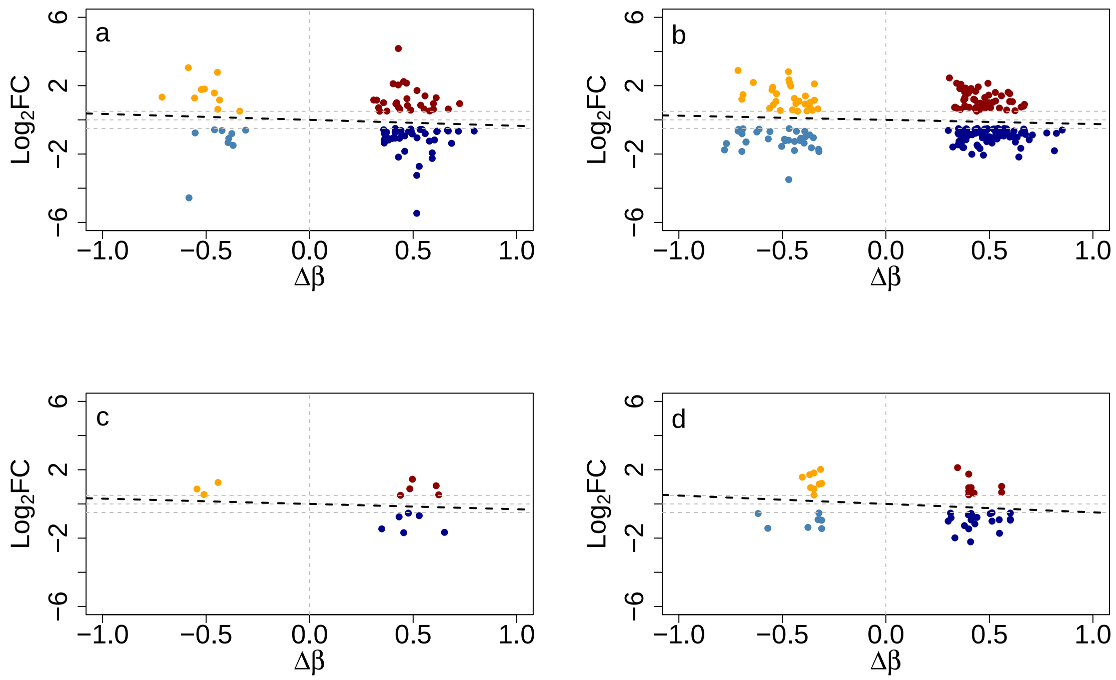


Figure 14: Correlation between $Log_2FC$ and $\Delta\beta$ values in genomic elements that do not overlap CpG Islands. Panels show the correlation between $Log_2FC$ and $\Delta\beta$ values of DEG with differentially methylated regulatory element (a), gene body (b), 3'UTR (c) and enhancer (d). All the DMRs taken into consideration in this analysis do not overlap with a CpG island. Dotted line represent the fitted linear regression curve.

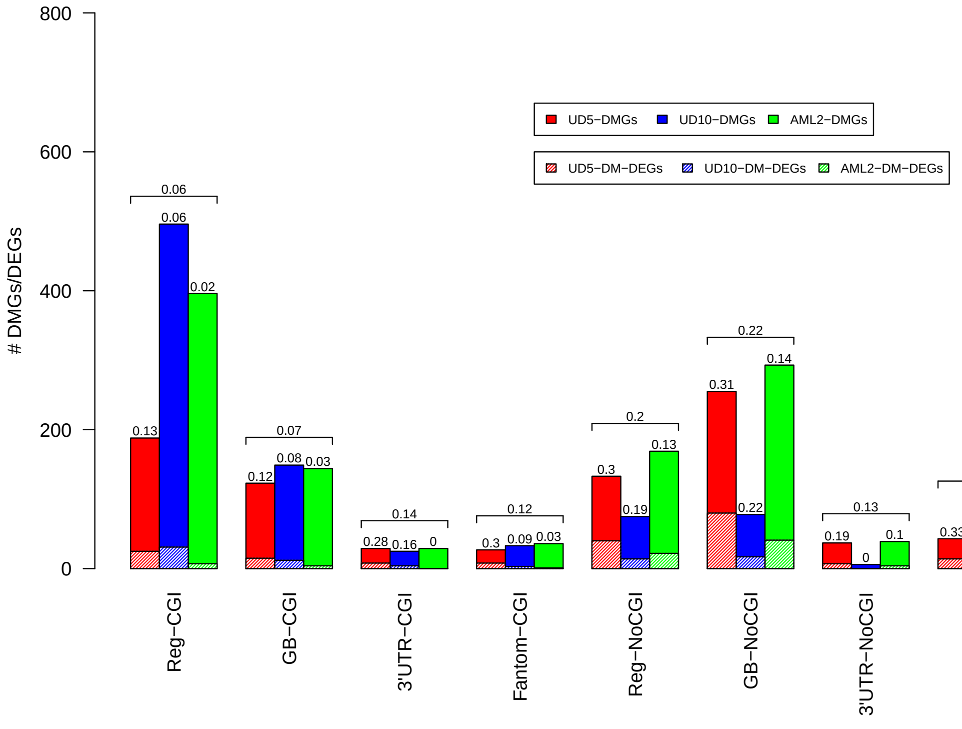


Figure 15: Hyper-methylated genes and differentially expressed genes on the three AML pairs. Barplots report the total number of genes hyper-methylated (solid bars) genes and the numbers of hyper-methylated genes that are also DEGs (textured bars) for each genic element (regulatory element (Reg), gene body (GB), 3'UTR and Fantom) in CGIs and in sparse CpG regions (No-CGI). Numbers above bars show the percentage of Hyper-methylated genes that are also DEGs. Horizontal brackets above each group of three bars summarize average percentages within the three samples.

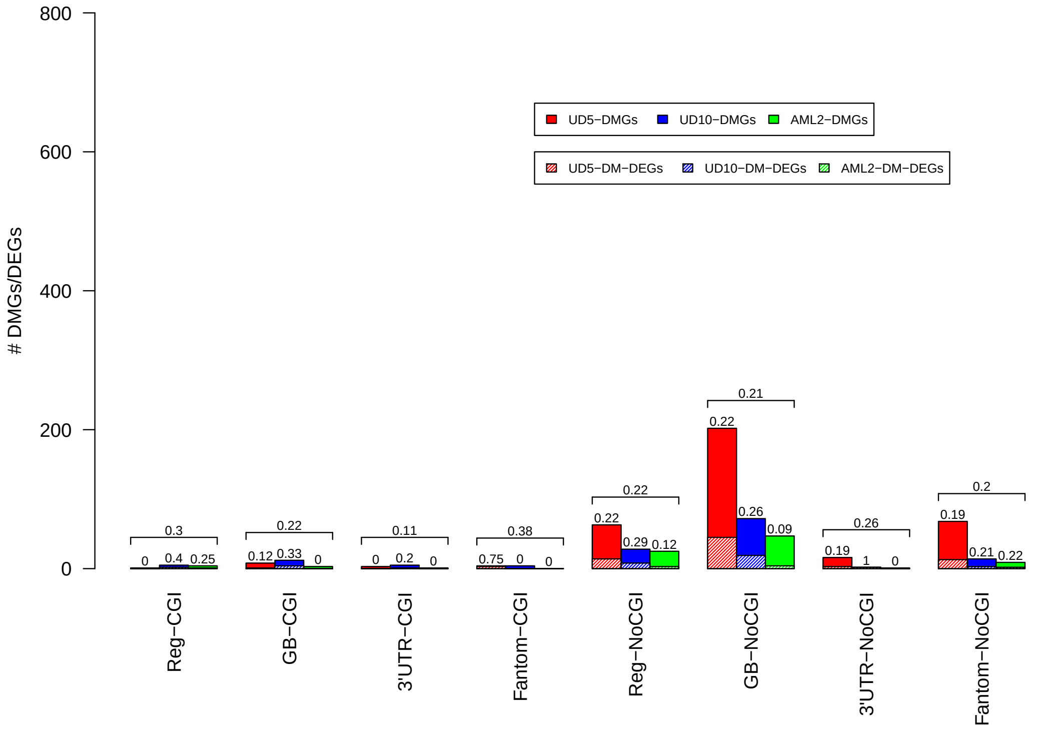


Figure 16: Hypo-methylated genes and differentially expressed genes on the three AML pairs. Barplots report the total number of hypo-methylated (solid bars) genes and the numbers of hypo-methylated genes that are also DEGs (textured bars) for each genic element (regulatory element (Reg), gene body (GB), 3'UTR and Fantom) in CGIs and in sparse CpG regions (No-CGI). Numbers above bars show the percentage of hypo-methylated genes that are also DEGs. Horizontal brackets above each group of three bars summarize average percentages within the three samples.

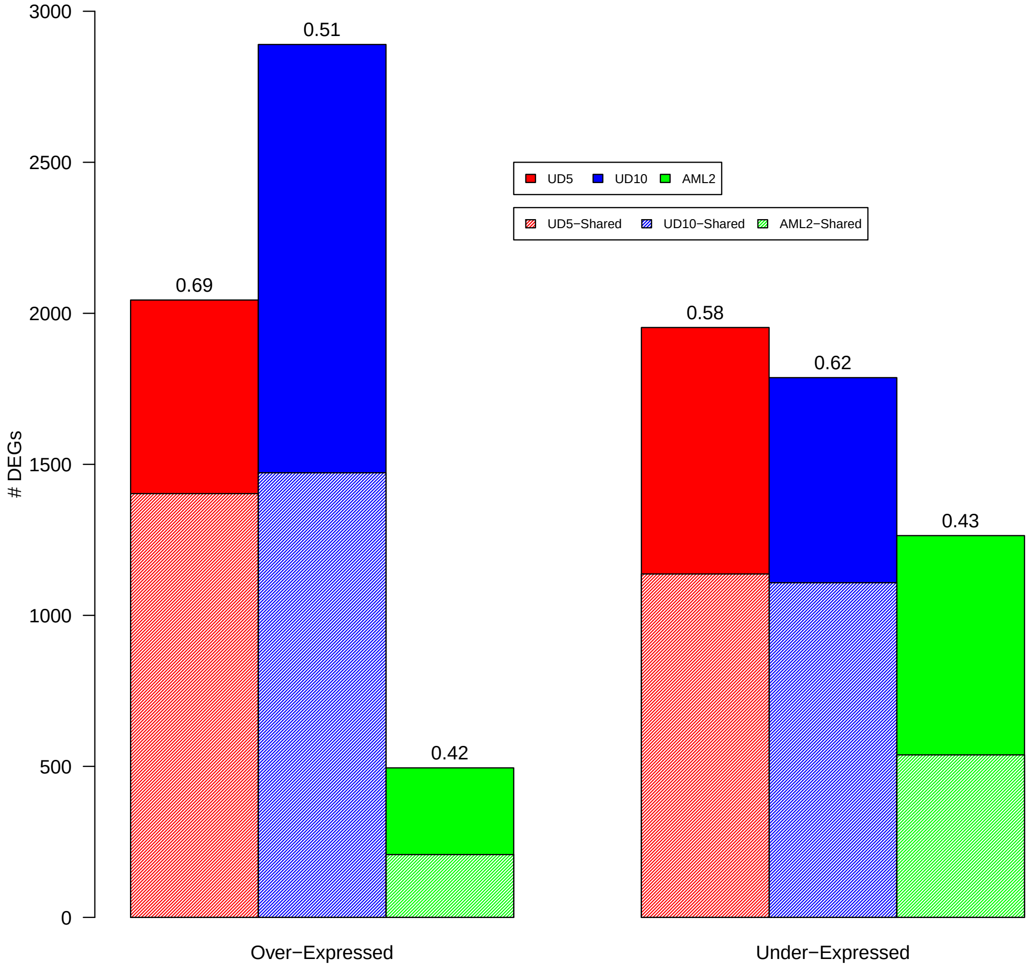


Figure 17: Differentially expressed (DEGs) identified by DESeq2 in the three AML pairs. Barplots report the total number of over-expressed (left) and under-expressed (right) genes detected by DESeq2 (adjusted p-value$< 0.05$ and absolute log2FC $> 0.5$) in the three AML pairs. Textured bars show the number of DEGs shared with the other two sample. Numbers above bars show the percentage of DEGs that are shared with the other two sample.

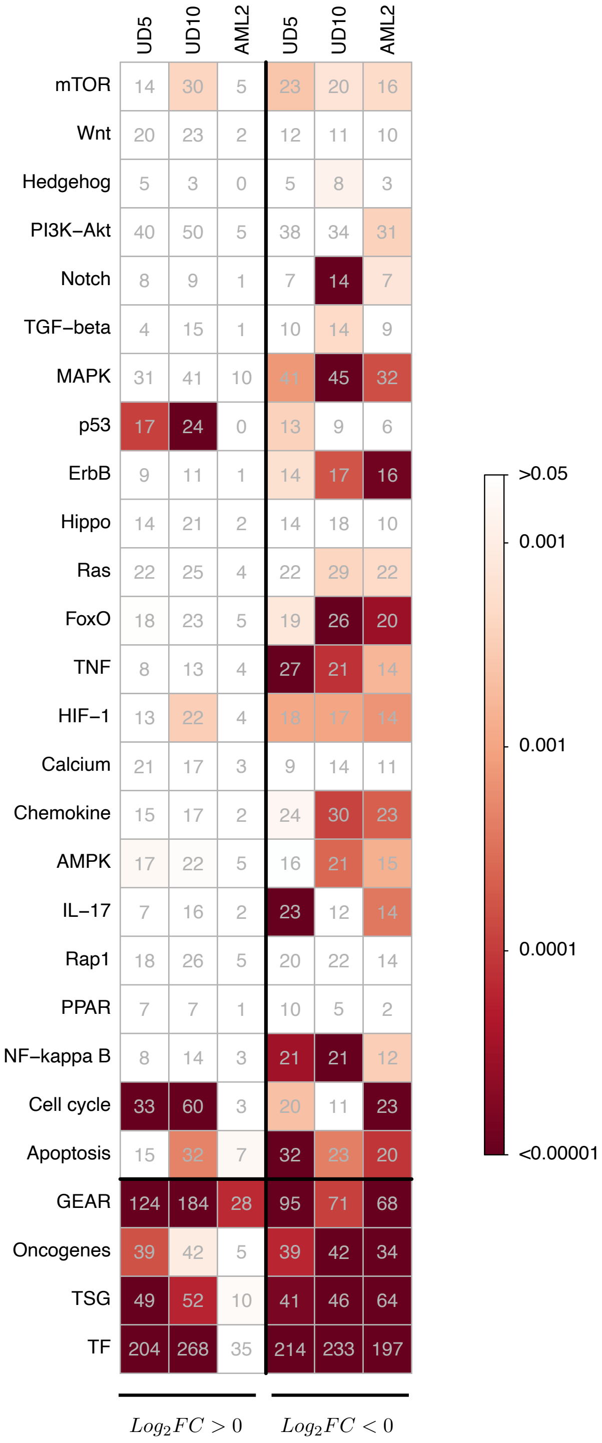


Figure 18: Over-representation analysis of DE genes. Each column of the matrix reports the results of over-representation analysis for the over- and under-expressed genes of each of the three sample. Fisher exact test is calculated for TF, COSMIC, NCG and for cancer-related pathways selected from KEGG and Reactome databases

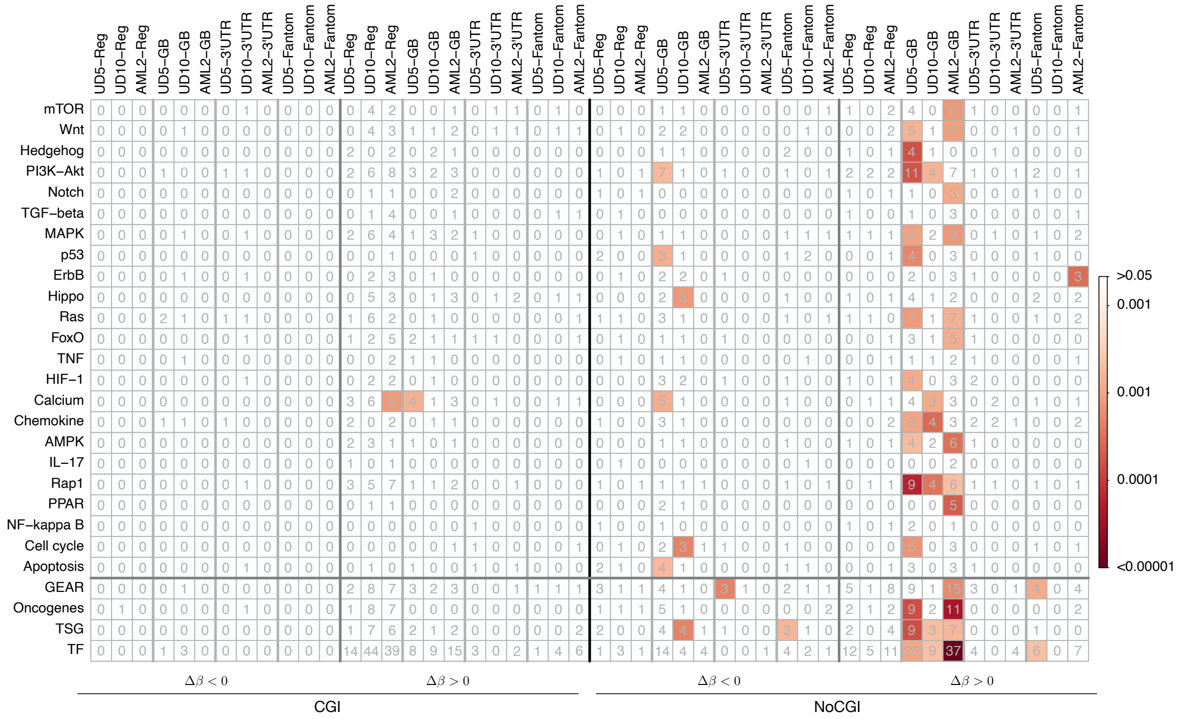


Figure 19: results of ORA of genes with DMR affecting 5' regulatory region (Reg), the gene body (GB), the 3'UTR and the enhancer region predicted by FANTOM. The analysis is performed separately for DMRs overlapping CpG islands (CGI) and outside CpG islands (NoCGI) for cancer-related pathways selected from KEGG database, TF, TSG and Oncogenes selected by COSMIC and GEAR genes. In each cell is reported the total number of DM genes for each category, while color reflect the p-value of the ORA analysis. P-values are obtained by testing (with Fisher exact test) the number of genes that overlap DMRs.

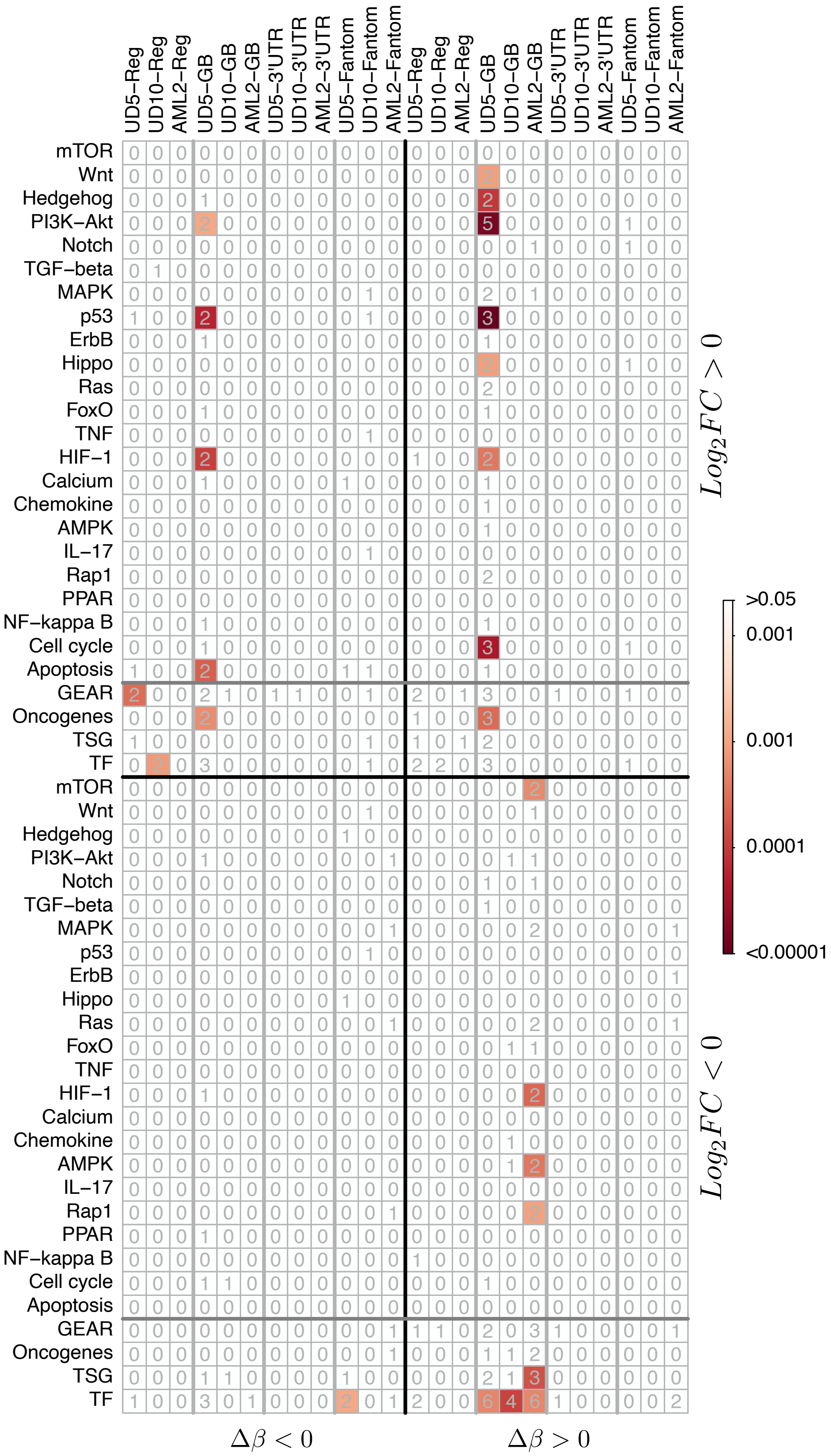


Figure 20: Over-representation analysis of DM-DE genes with DMRs overlapping sparse CpG regions at different genomic elements. Each cell of the matrix reports the results of overrepresentation analysis for all the four epigenetic regulation (hypo-methylation/over-expression, hyper-methylation/under-expression, hyper-methylation/over-expression and hypo-methylation/under-expression). The analysis was performed separately for genes affected by DMRs at the 5' regulatory region (Reg), the gene body (GB), the 3'UTR and at the enhancer region predicted by FANTOM. The numbers in each cell represent the total number of genes for each category, while the color intensity reflects statistical significance according to colorbar. Fisher exact test and number of genes were calculated for TF, TSG, Oncogenes, GEAR and for cancer-related pathways selected from KEGG.

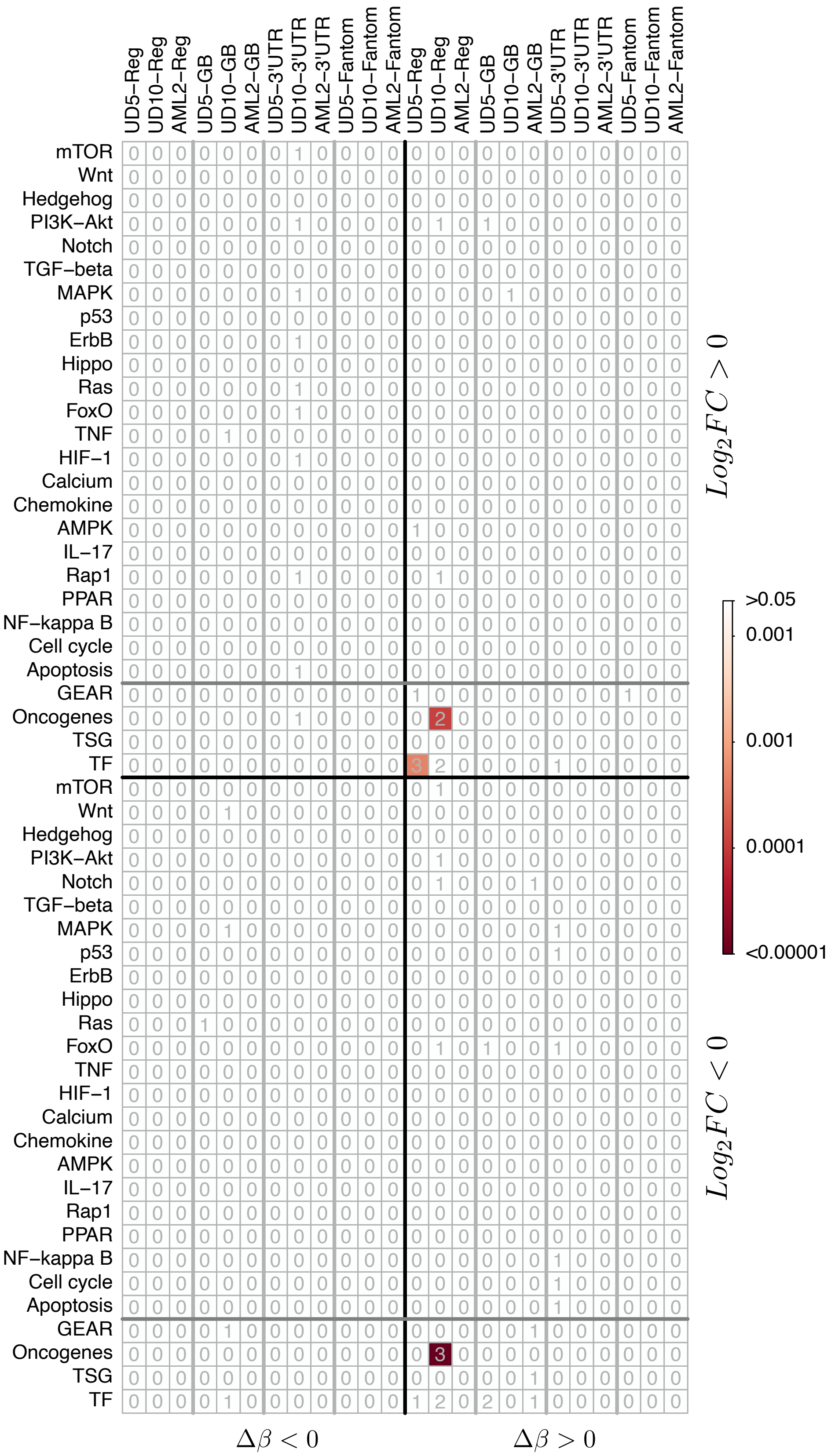


Figure 21: Over-representation analysis of DM-DE genes with DMRs overlapping CGIs at different genomic elements. Each cell of the matrix reports the results of overrepresentation analysis for all the four epigenetic regulation (hypo-methylation/over-expression, hyper-methylation/under-expression, hyper-methylation/over-expression and hypo-methylation/under-expression). The analysis was performed separately for genes affected by DMRs at the 5' regulatory region (Reg), the gene body (GB), the 3'UTR and at the enhancer region predicted by FANTOM. The numbers in each cell represent the total number of genes for each category, while the color intensity reflects statistical significance according to colorbar. Fisher exact test and number of genes were calculated for TF, TSG, Oncogenes, GEAR and for cancer-related pathways selected from KEGG.

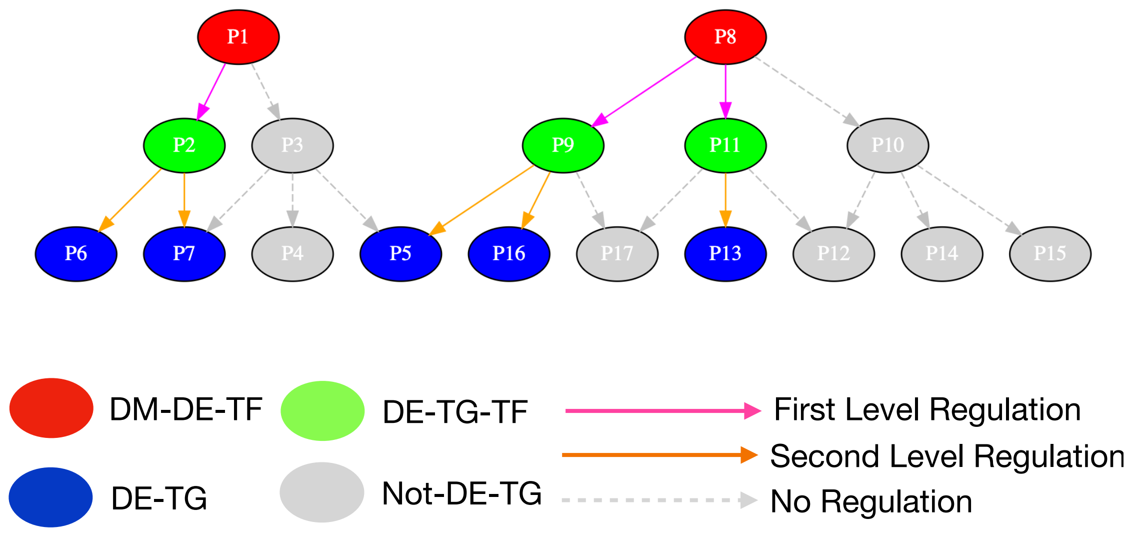


Figure 22: Figure shows the scheme we used to study the effect of DMRs on gene regulatory network cascade. Red nodes represent differentially methylated and differentially expressed TFs (DM-DE-TF). Green nodes are differentially expressed TFs regulated by DM-DE-TF (DE-TG-TF). Blue nodes are differentially expressed genes regulated by DM-DE-TF or by DE-TG-TF. Grey nodes are target genes that are not differentially expressed (Not-DE-TG). Magenta arrows indicate a first level regulation (DM-DE-TF regulate a DE target gene). Orange arrow indicate a second level regulation (DE-TG-TF regulate a DE target gene). Grey arrows indicate no regulation.

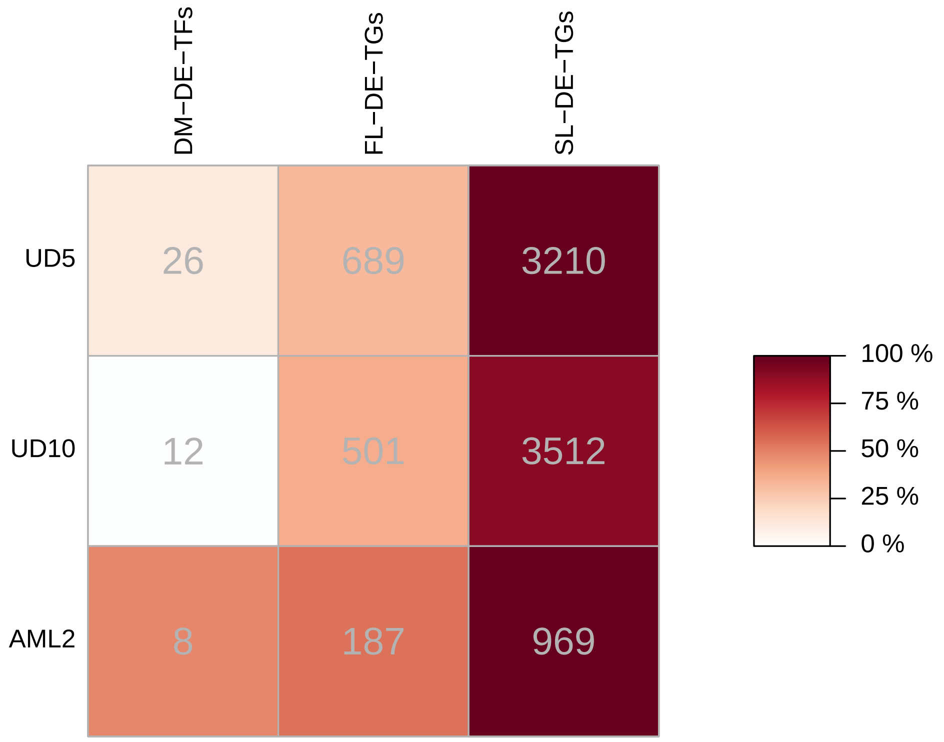


Figure 23: The table summarizes the total number of DEG in each level of the regulatory cascade induced by DMRs. In each cell of the first column is reported the total number of differentially methylated and differentially expressed TFs (DM-DE-TS). In the second column the total number of first-level target genes (FL-DE-TGs) and in the third column the total number of second-level target genes (SL-DE-TGs). For each AML sample the color intensity in each cell shows the proportion of genes shared with the other two samples.

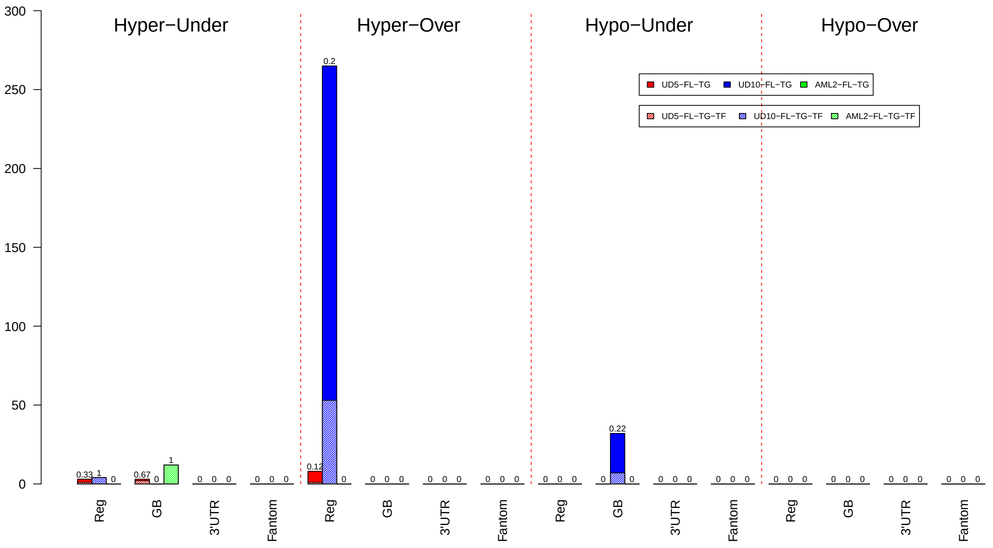


Figure 24: The barplot reports the total number of first-level target genes (FL-DE-TGs) potentially regulated by DM-DE-TFs with DMRs in CGIs. Each bar reports the number of FL-DE-TGs regulated by under- or over-expressed DM-DE-TFs with hyper- or hypo-methylated DMRs at 5' regulatory-elements, gene bodies, 3'UTRs and enhancers. For each category, textured bars show the number of FL-DE-TGs shared with the other two samples. Numbers above bars show the percentage of DEGs that are shared with the other two sample.

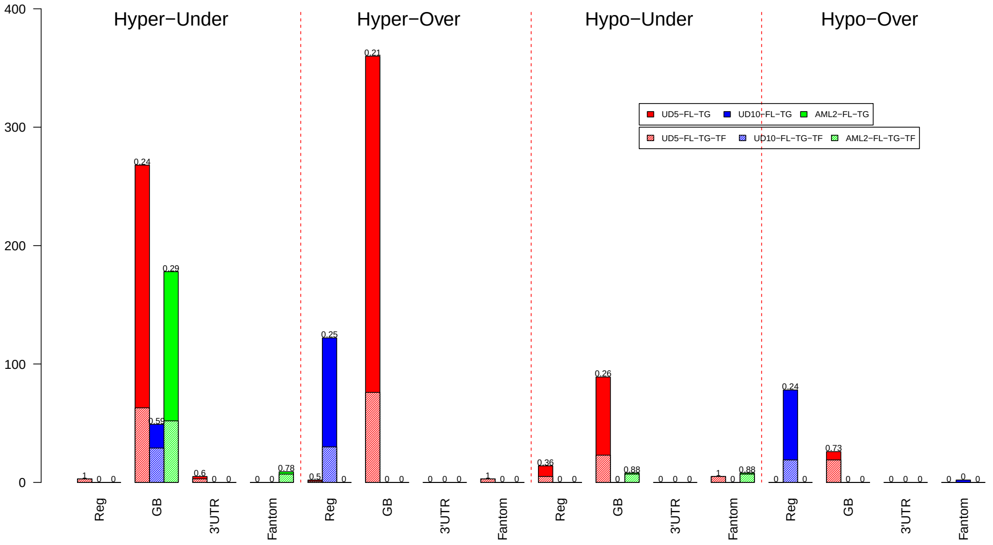


Figure 25: The barplot reports the total number of first-level target genes (FL-DE-TGs) potentially regulated by DM-DE-TFs with DMRs in sparse CpG regions. Each bar reports the number of FL-DE-TGs regulated by under- or over-expressed DM-DE-TFs with hyper- or hypo-methylated DMRs at 5' regulatory-elements, gene bodies, 3'UTRs and enhancers. For each category, textured bars show the number of FL-DE-TGs shared with the other two samples. Numbers above bars show the percentage of DEGs that are shared with the other two sample.

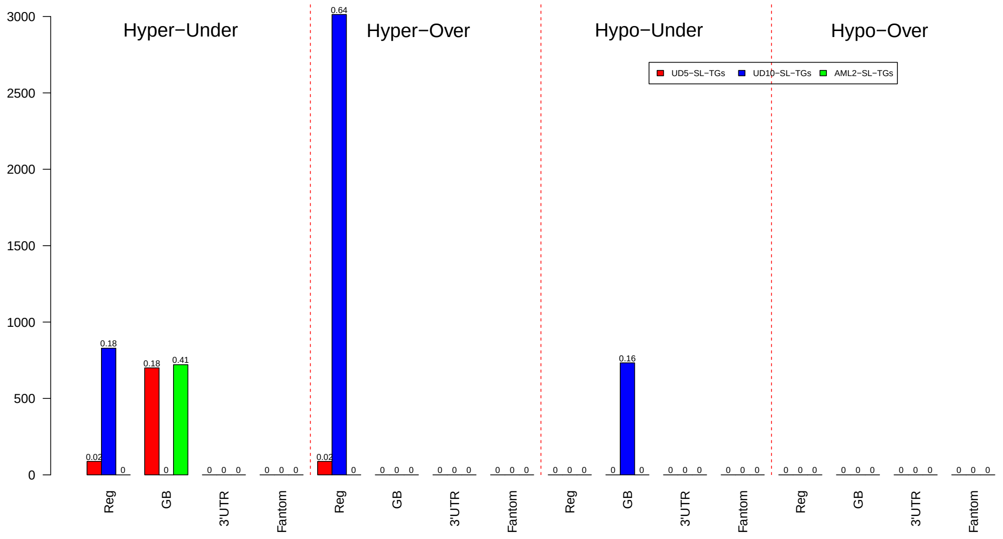


Figure 26: The barplot reports the total number of first- and second-level target genes (FL-DE-TGs) potentially regulated by DM-DE-TFs with DMRs in CGIs. Each bar reports the number of FL-and SL-DE-TGs regulated by under- or over-expressed DM-DE-TFs with hyper- or hypo-methylated DMRs at 5' regulatory-elements, gene bodies, 3'UTRs and enhancers. For each category, numbers above bars show the fraction of FL-and SL-DE-TGs with respect to all under- or over-expressed genes for each sample.

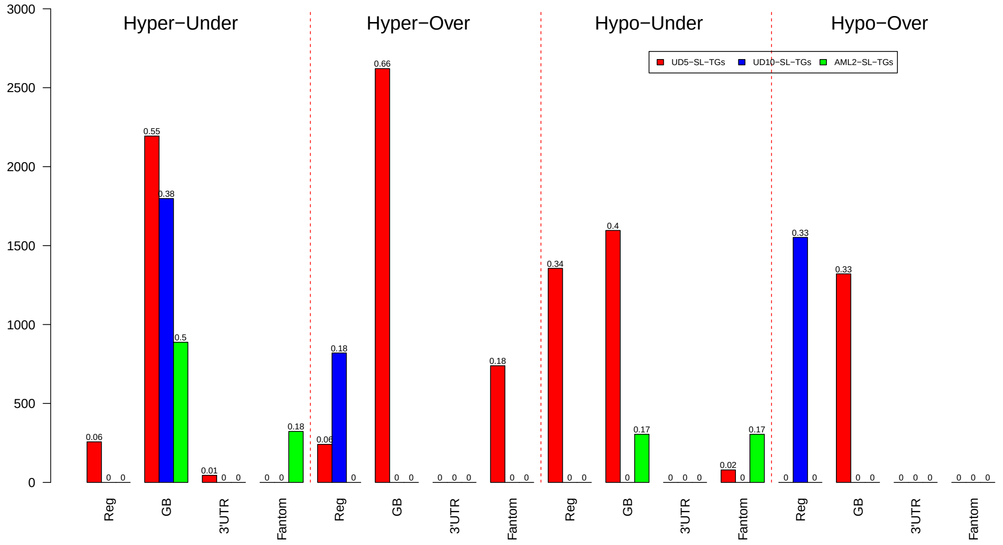


Figure 27: The barplot reports the total number of first- and second-level target genes (FL-DE-TGs) potentially regulated by DM-DE-TFs with DMRs in sparse CpG regions. Each bar reports the number of FL- and SL-DE-TGs regulated by under- or over-expressed DM-DE-TFs with hyper- or hypo-methylated DMRs at 5' regulatory-elements, gene bodies, 3'UTRs and enhancers. For each category, numbers above bars show the fraction of FL-and SL-DE-TGs with respect to all under- or over-expressed genes for each sample.

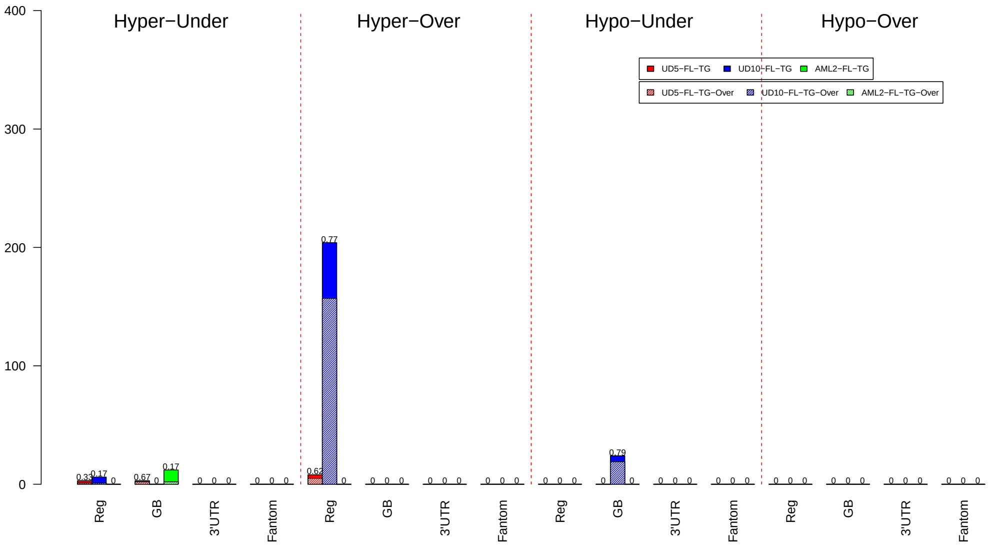


Figure 28: The barplot reports the total number of over- and under-expressed first-level target genes (FL-DE-TGs) potentially regulated by DM-DE-TFs with DMRs in CGIs. Each bar reports the number of FL-DE-TGs regulated by under- or over-expressed DM-DE-TFs with hyper- or hypo-methylated DMRs at 5' regulatory-elements, gene bodies, 3'UTRs and enhancers. For each category, textured bars show the number of over-expressed FL-DE-TGs while solid bars the number of under-expressed FL-DE-TGs. Numbers above bars show the percentage of over-expressed FL-DE-TGs.

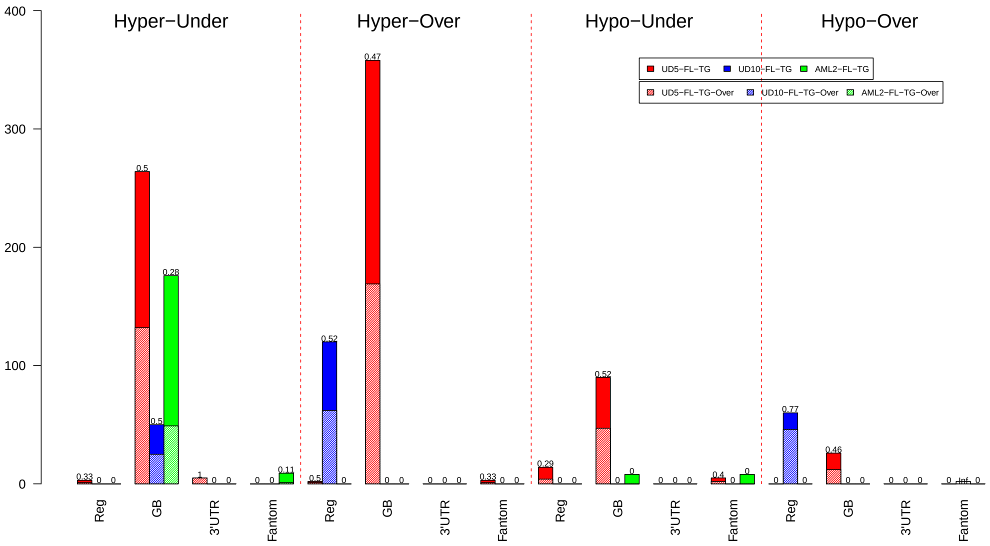


Figure 29: The barplot reports the total number of over- and under-expressed second-level target genes (FL-DE-TGs) potentially regulated by DM-DE-TFs with DMRs in sparse CpG regions. Each bar reports the number of FL-DE-TGs regulated by under- or over-expressed DM-DE-TFs with hyper- or hypo-methylated DMRs at 5' regulatory-elements, gene bodies, 3'UTRs and enhancers. For each category, textured bars show the number of over-expressed FL-DE-TGs while solid bars the number of under-expressed FL-DE-TGs. Numbers above bars show the percentage of over-expressed FL-DE-TGs.

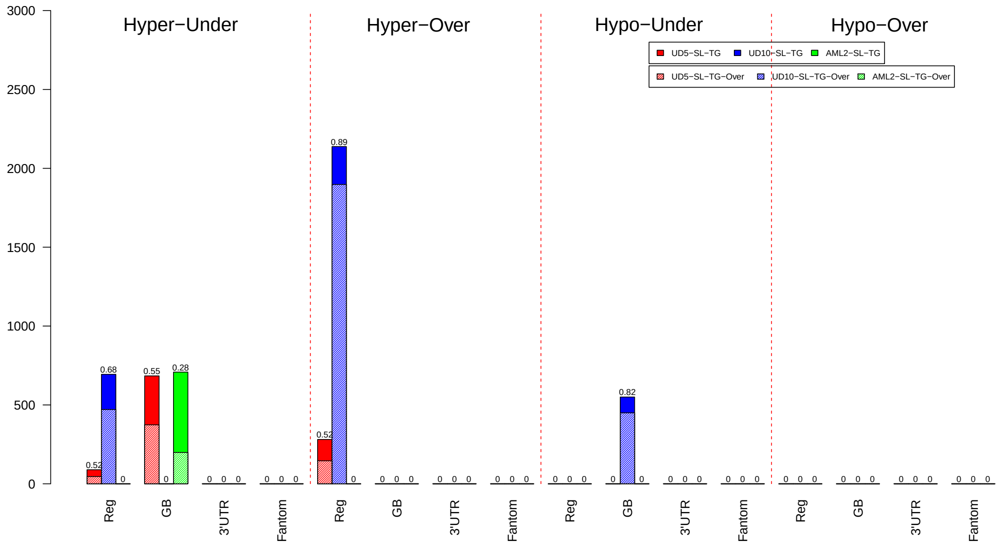


Figure 30: The barplot reports the total number of over- and under-expressed second-level target genes (SL-DE-TGs) potentially regulated by DM-DE-TFs with DMRs in CGIs. Each bar reports the number of SL-DE-TGs regulated by under- or over-expressed DM-DE-TFs with hyper- or hypo-methylated DMRs at 5' regulatory-elements, gene bodies, 3'UTRs and enhancers. For each category, textured bars show the number of over-expressed SL-DE-TGs while solid bars the number of under-expressed SL-DE-TGs. Numbers above bars show the percentage of over-expressed SL-DE-TGs.

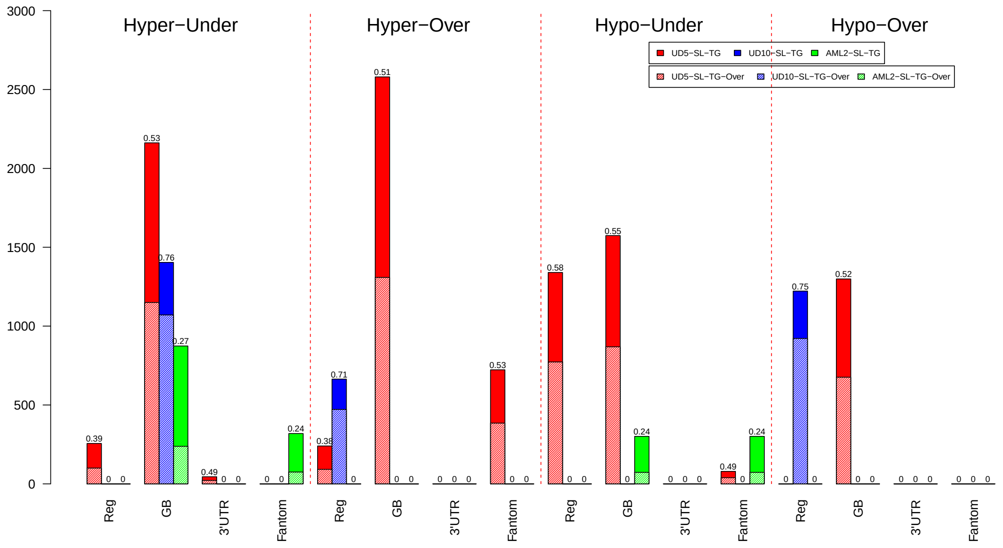


Figure 31: The barplot reports the total number of over- and under-expressed second-level target genes (SL-DE-TGs) potentially regulated by DM-DE-TFs with DMRs in sparse CpG regions. Each bar reports the number of SL-DE-TGs regulated by under- or over-expressed DM-DE-TFs with hyper- or hypo-methylated DMRs at 5' regulatory-elements, gene bodies, 3'UTRs and enhancers. For each category, textured bars show the number of over-expressed SL-DE-TGs while solid bars the number of under-expressed SL-DE-TGs. Numbers above bars show the percentage of over-expressed SL-DE-TGs.

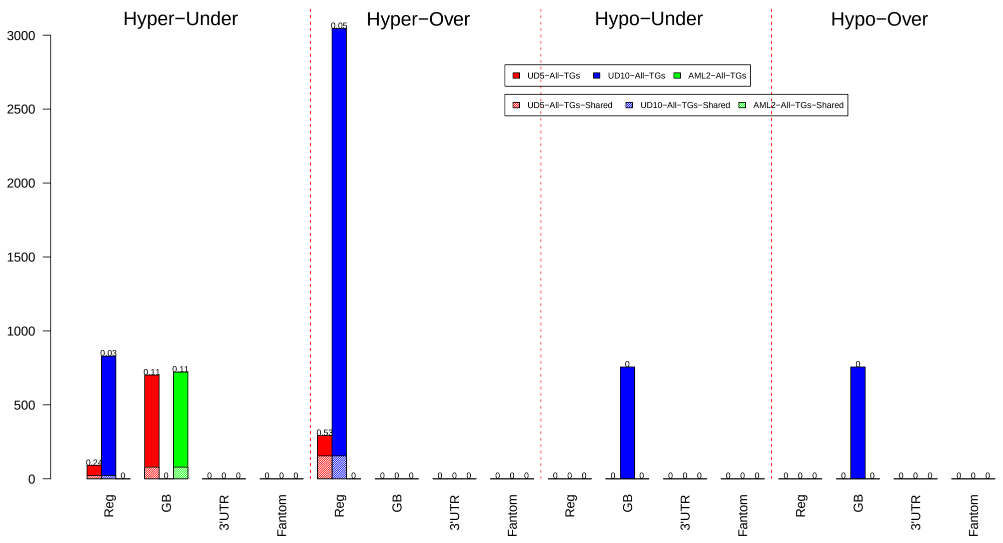


Figure 32: The barplot reports the total number of first- and second-level target genes (FL-DE-TGs) potentially regulated by DM-DE-TFs with DMRs in CGIs. Each bar reports the number of FL- and SL-DE-TGs regulated by under- or over-expressed DM-DE-TFs with hyper- or hypo-methylated DMRs at 5' regulatory-elements, gene bodies, 3'UTRs and enhancers. For each category, textured bars show the number of FL- and SL-DE-TGs shared with the other two samples. Numbers above bars show the percentage of DEGs that are shared with the other two sample.

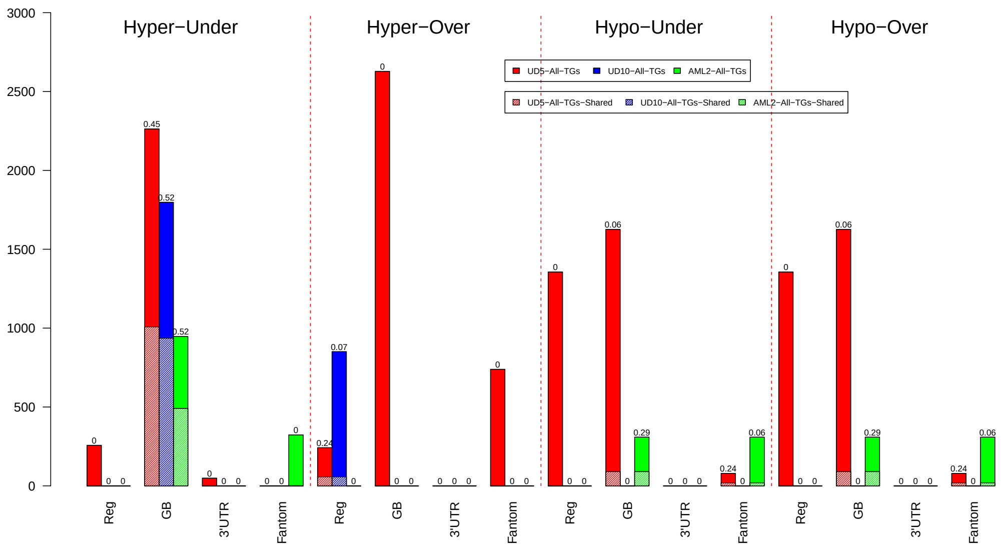


Figure 33: The barplot reports the total number of first- and second-level target genes (FL-DE-TGs) potentially regulated by DM-DE-TFs with DMRs in sparse CpG regions. Each bar reports the number of FL- and SL-DE-TGs regulated by under- or over-expressed DM-DE-TFs with hyper- or hypo-methylated DMRs at 5' regulatory-elements, gene bodies, 3'UTRs and enhancers. For each category, textured bars show the number of FL- and SL-DE-TGs shared with the other two samples. Numbers above bars show the percentage of DEGs that are shared with the other two sample.

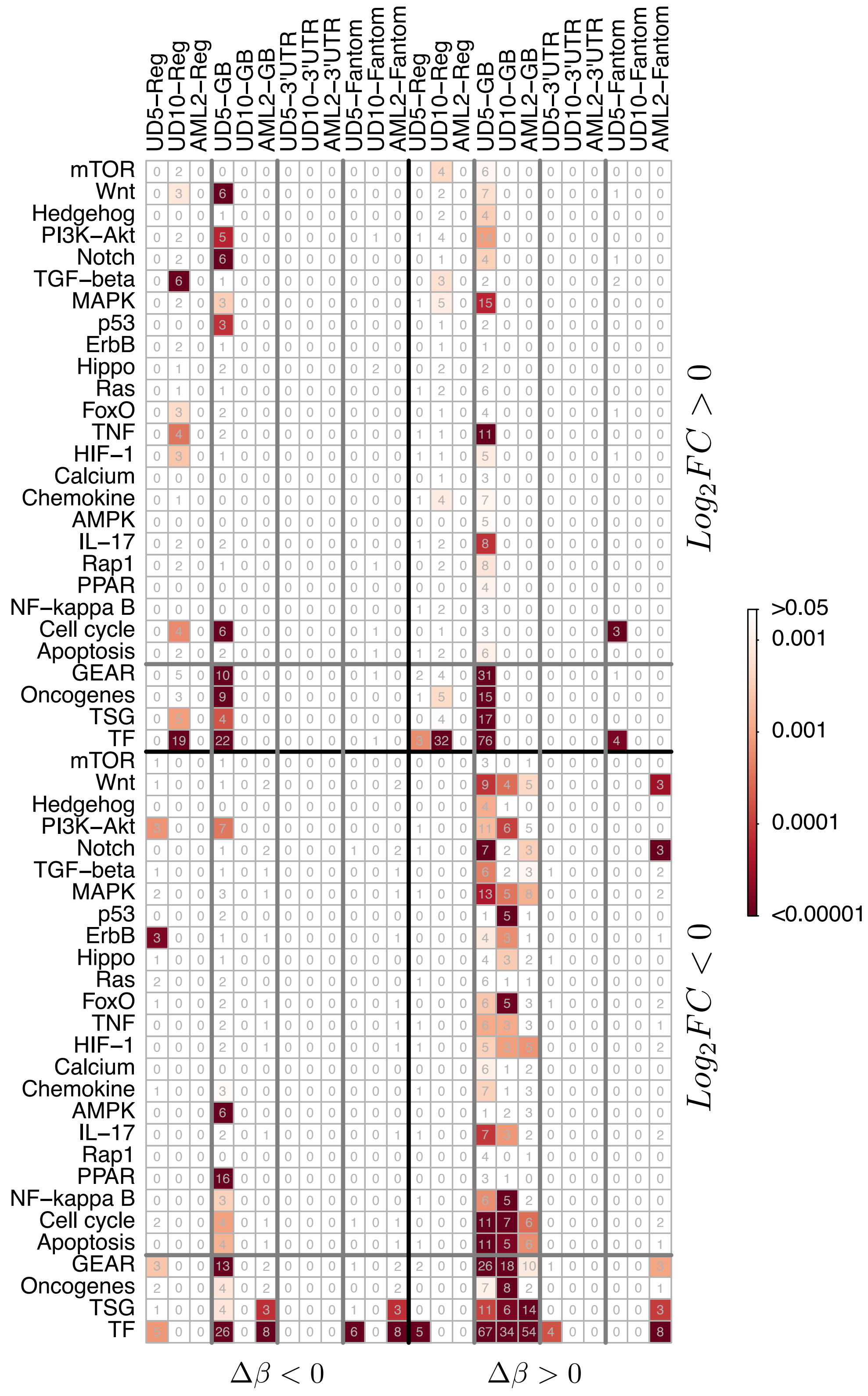


Figure 34: Figure reports the results of ORA on the first-level regulatory network cascade induced by DMRs at sparse CpG regions. Each cell of the matrix reports the results of overrepresentation analysis for all the four epigenetic regulation (hypo-methylation/over-expression, hyper-methylation/under-expression, hyper-methylation/over-expression and hypo-methylation/under-expression). The analysis was performed separately for genes affected by DMRs at the 5' regulatory region (Reg), the gene body (GB), the 3'UTR and at the enhancer region predicted by FANTOM. The numbers in each cell represent the total number of genes for each category, while the color intensity reflects statistical significance according to colorbar. Fisher exact test and number of genes were calculated for cancer-related pathways selected from KEGG database, TFs, TSG and Oncogenes selected by COSMIC and GEAR genes.

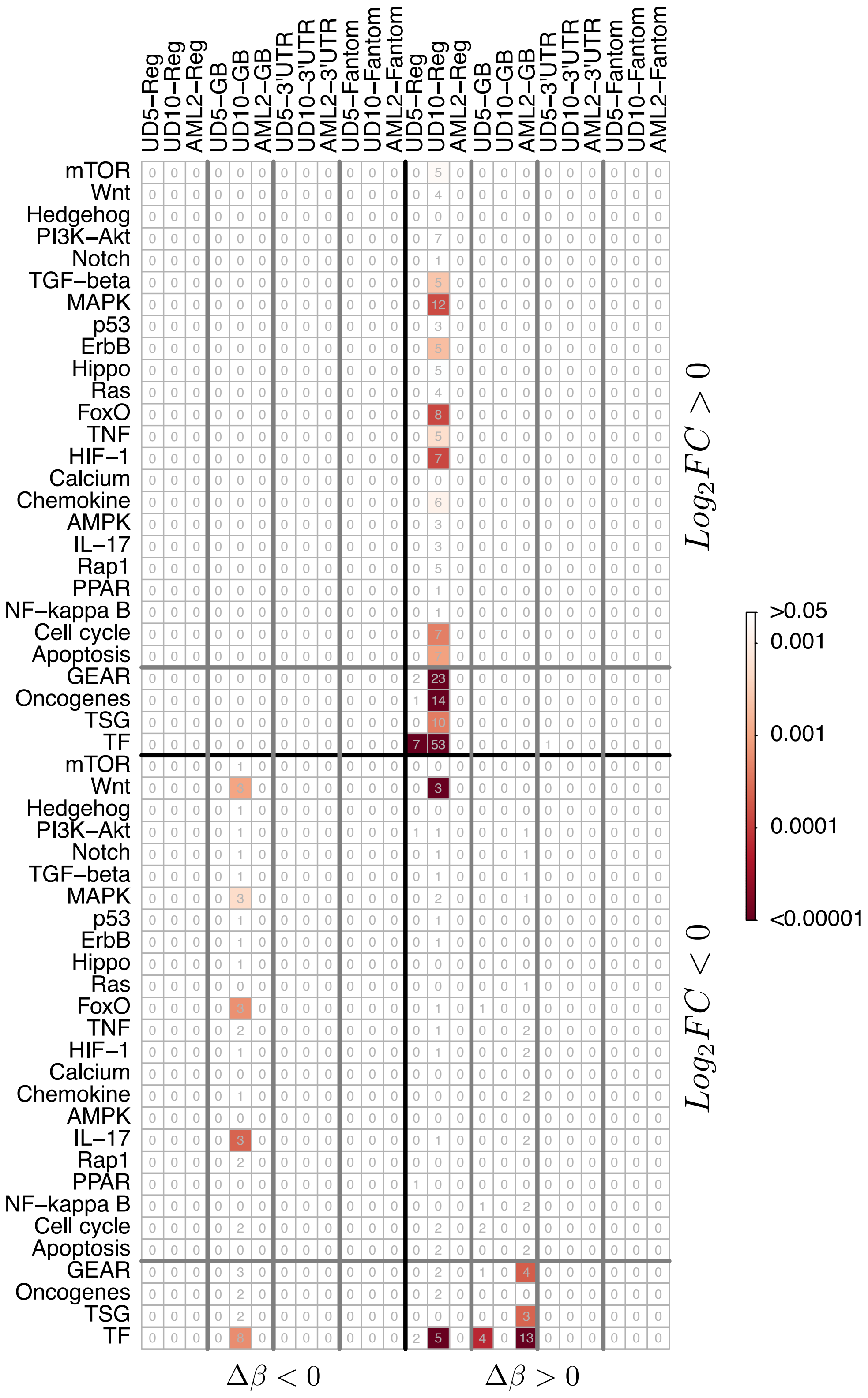


Figure 35: Figure reports the results of ORA on the first-level regulatory network cascade induced by DMRs at CGIs. Each cell of the matrix reports the results of overrepresentation analysis for all the four epigenetic regulation (hypo-methylation/over-expression, hyper-methylation/under-expression, hyper-methylation/over-expression and hypo-methylation/under-expression). The analysis was performed separately for genes affected by DMRs at the 5' regulatory region (Reg), the gene body (GB), the 3'UTR and at the enhancer region predicted by FANTOM. The numbers in each cell represent the total number of genes for each category, while the color intensity reflects statistical significance according to colorbar. Fisher exact test and number of genes were calculated for cancer-related pathways selected from KEGG database, TFs, TSG and Oncogenes selected by COSMIC and GEAR genes.

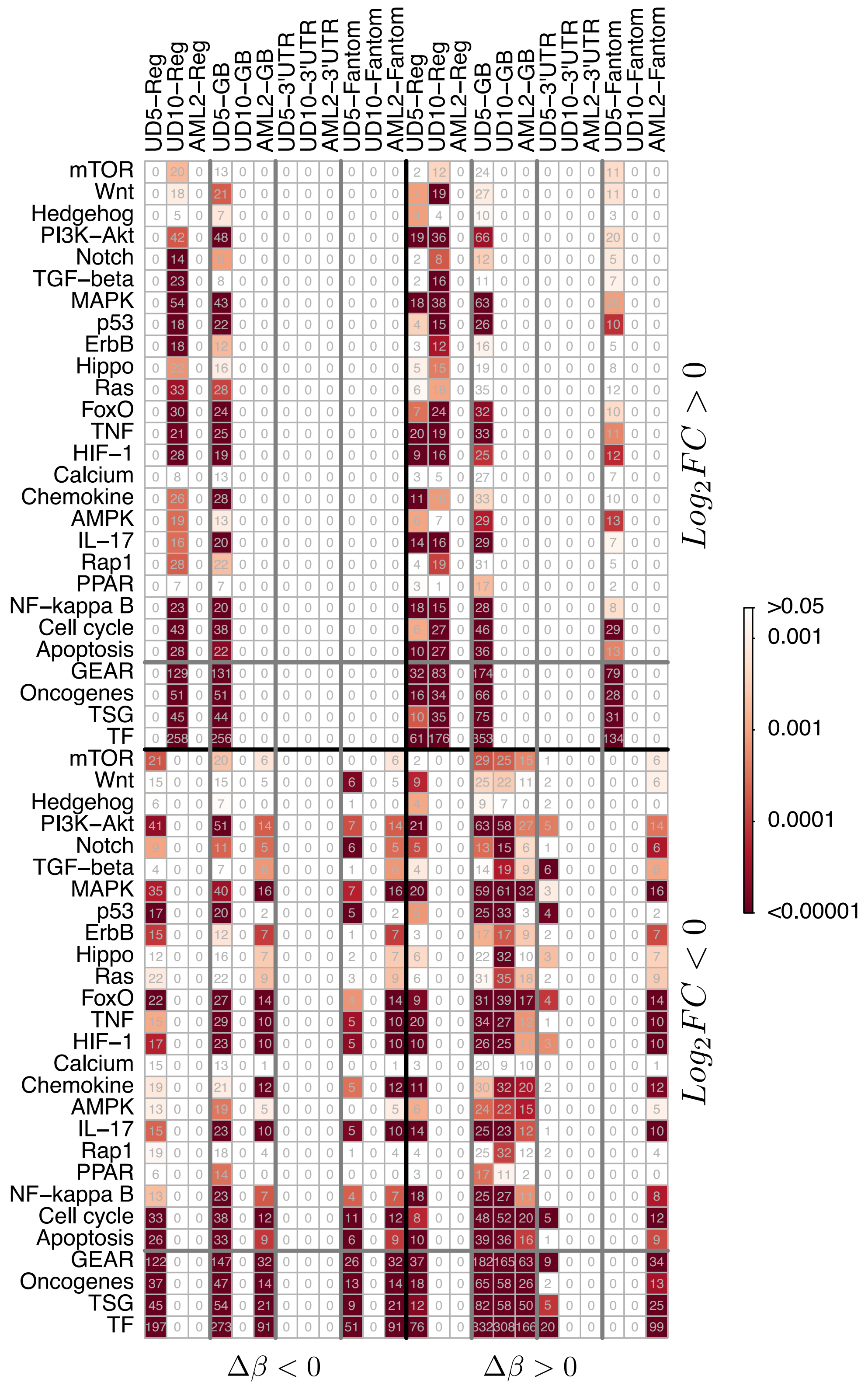


Figure 36: Figure reports the results of ORA on the second-level regulatory network cascade induced by DMRs at sparse CpG regions. Each cell of the matrix reports the results of overrepresentation analysis for all the four epigenetic regulation (hypo-methylation/over-expression, hyper-methylation/under-expression, hyper-methylation/over-expression and hypo-methylation/under-expression). The analysis was performed separately for genes affected by DMRs at the 5' regulatory region (Reg), the gene body (GB), the 3'UTR and at the enhancer region predicted by FANTOM. The numbers in each cell represent the total number of genes for each category, while the color intensity reflects statistical significance according to colorbar. Fisher exact test and number of genes were calculated for cancer-related pathways selected from KEGG database, TFs, TSG and Oncogenes selected by COSMIC and GEAR genes.

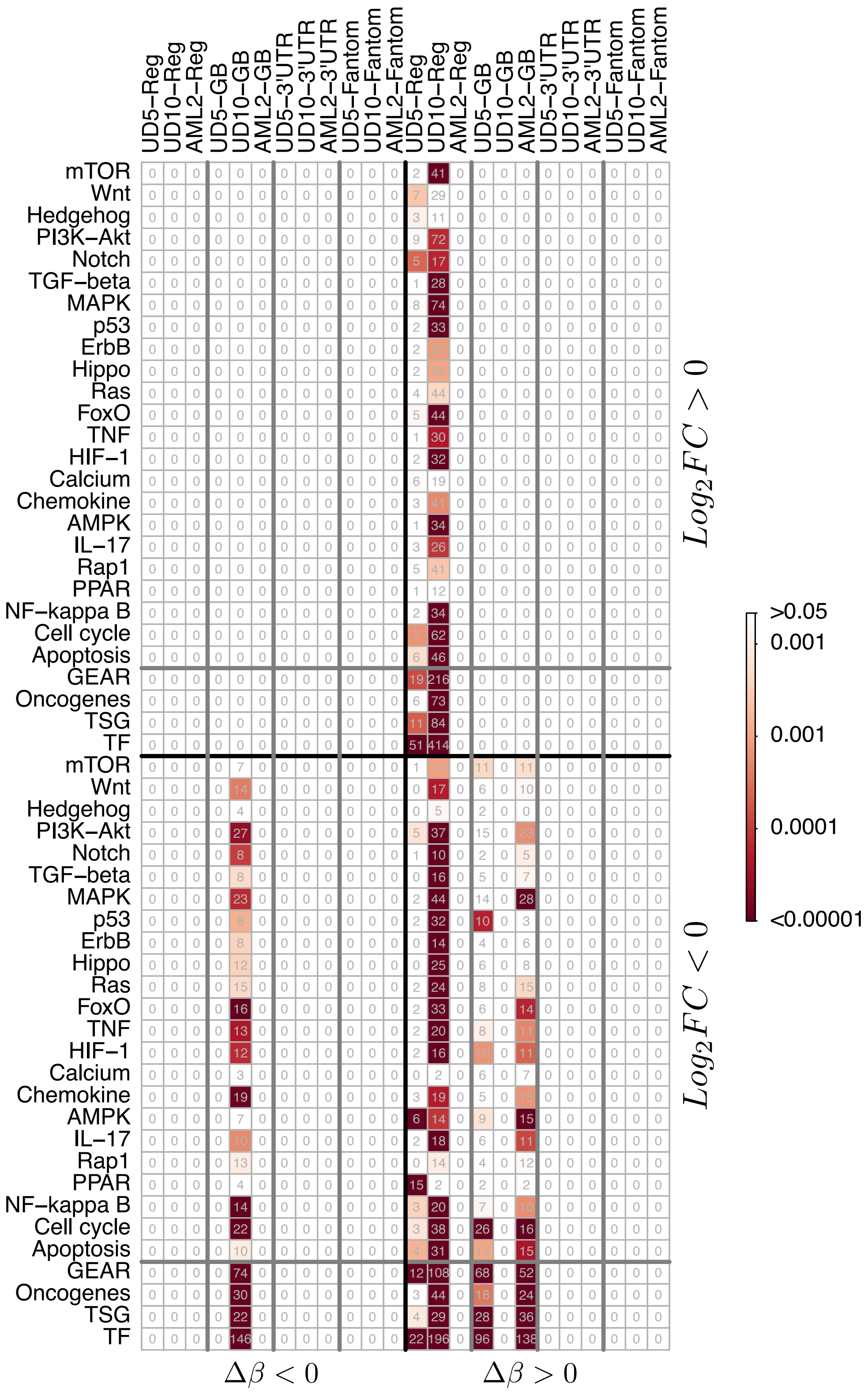


Figure 37: Figure reports the results of ORA on the second-level regulatory network cascade induced by DMRs at CGIs. Each cell of the matrix reports the results of overrepresentation analysis for all the four epigenetic regulation (hypo-methylation/over-expression, hyper-methylation/under-expression, hyper-methylation/over-expression and hypo-methylation/under-expression). The analysis was performed separately for genes affected by DMRs at the 5' regulatory region (Reg), the gene body (GB), the 3'UTR and at the enhancer region predicted by FANTOM. The numbers in each cell represent the total number of genes for each category, while the color intensity reflects statistical significance according to colorbar. Fisher exact test and number of genes were calculated for cancer-related pathways selected from KEGG database, TFs, TSG and Oncogenes selected by COSMIC and GEAR genes.

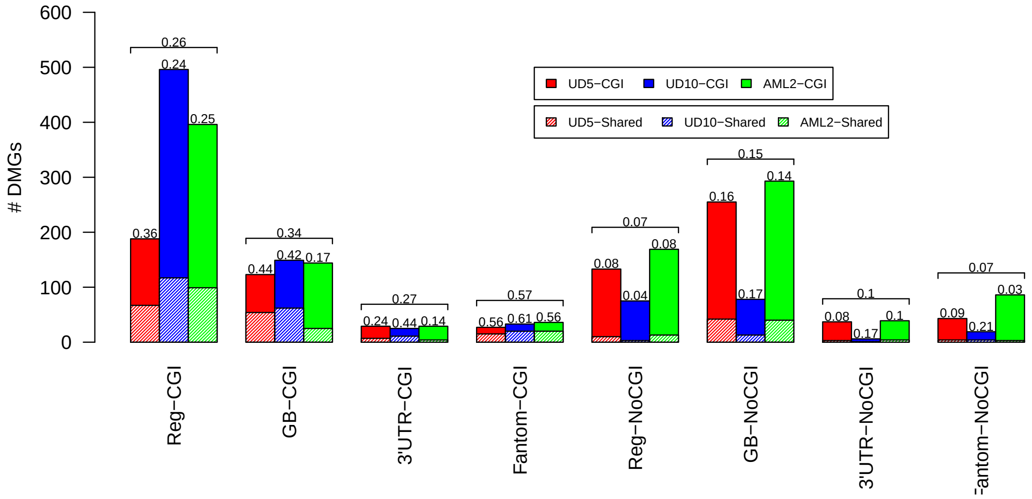


Figure 38: The barplot reports the number of hyper-methylated DMGs shared by each AML sample with the other two samples. Solid bars report the total number of hyper-methylated DMGs, while textured bars the number of shared DMGs. Numbers are reported for DMGs with DMRs at 5' regulatory-elements, gene bodies, 3'UTRs and enhancers (Fantom) overlapping CGIs and sparse CpG regions (NoCGI). Numbers above bars show the percentage of DMGs that are shared with the other two sample. Horizontal brackets above each group of three bars summarize average percentages within the three samples.

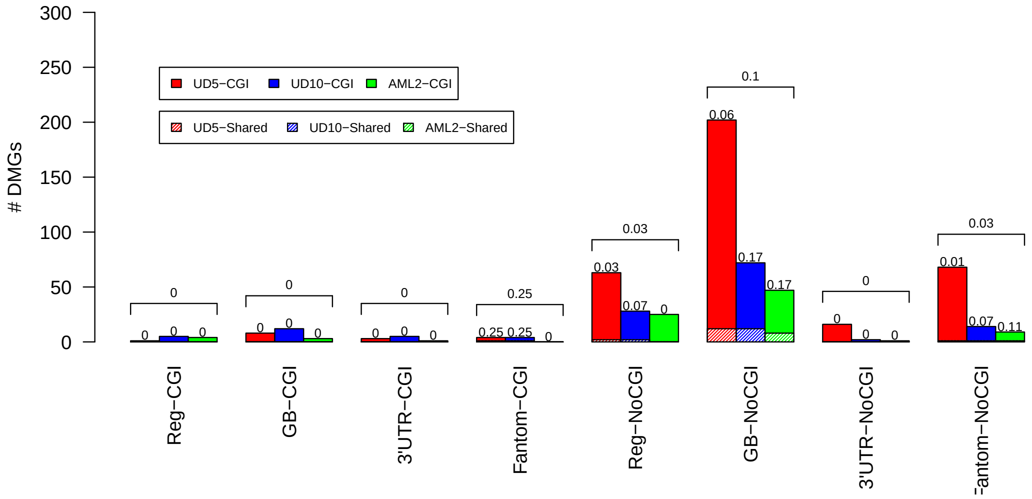


Figure 39: The barplot reports the number of hypo-methylated DMGs shared by each AML sample with the other two samples. Solid bars report the total number of hypo-methylated DMGs, while textured bars the number of shared DMGs. Numbers are reported for DMGs with DMRs at 5' regulatory-elements, gene bodies, 3'UTRs and enhancers (Fantom) overlapping CGIs and sparse CpG regions (NoCGI). Numbers above bars show the percentage of DMGs that are shared with the other two sample. Horizontal brackets above each group of three bars summarize average percentages within the three samples.
